# Targeting integrin alpha5 receptor in pancreatic stellate cells to diminish tumor-promoting effects in pancreatic cancer

**DOI:** 10.1101/350678

**Authors:** Praneeth R. Kuninty, Ruchi Bansal, Sanne W.L. De Geus, Jonas Schnittert, Joop van Baarlen, Gert Storm, Maarten F. Bijlsma, Hanneke W. van Laarhoven, Peter J.K. Kuppen, Alexander L. Vahrmeijer, Arne Östman, Cornelis F.M. Sier, Jai Prakash

**Author notes:** Authors contributed equally (Ruchi Bansal, Sanne W.L. De Geus). Corresponding author’s Corresponding address: Section Targeted Therapeutics, Department of Biomaterials Science and Technology, Faculty of Science and Technology, University of Twente, Zuidhorst 254, 7500 AE Enschede, The Netherlands.

## Abstract

Pancreatic stellate cells (PSCs) are the main precursors of cancer-associated fibroblasts (CAFs) in pancreatic ductal adenocarcinoma (PDAC), known to induce cancer aggressiveness. Integrin alpha5 (ITGA5), a fibronectin receptor, was found to be overexpressed by CAFs in stroma and linked to poor overall survival (log-rank p=0.022, n=137) of patients with PDAC. *In vitro*, knockdown of ITGA5 in human PSCs (hPSCs) inhibited their adhesion, migration, and proliferation and also inhibited TGF-β-mediated differentiation. *In vivo*, co-injection of PANC-1 tumor cells and hPSCs (sh-ITGA5) developed tumors with reduced fibrosis and slower growth rate compared to those composed of PANC-1 and hPSC (sh-Ctrl). Furthermore, we developed a ITGA5-antagonizing peptidomimetic (AV3) which inhibited TGFβ-mediated hPSC differentiation by blocking ITGA5/FAK pathway. *In vivo*, treatment with AV3 intraperitoneally attenuated tumor fibrosis and thereby enhanced the efficacy of gemcitabine in patient-derived xenografts in mice. Altogether, this study reports the therapeutic importance of ITGA5 in PDAC and provides novel therapeutic peptidomimetic to enhance the effect of chemotherapy.

## Introduction

Pancreatic ductal adenocarcinoma (PDAC) is one of the most devastating cancers with a 5-year survival rate of less than 8% ^1^. Currently available therapies are insufficient to substantially halt the growth of PDAC, indicating an unmet clinical need to develop novel therapeutics. PDAC is characterized by abundant tumor stroma (up to 90% of the total tumor mass), which has been shown to promote tumor growth and metastasis, and confer resistance to chemotherapy, as well as act as a physical barrier by preventing tumor delivery of therapeutics ^2, 3, 4^. Pancreatic tumor stroma is composed of non-malignant cells such as cancer-associated fibroblasts (CAFs), immune cells, vasculature and a network of extracellular matrix (ECM), which interact with tumor cells in a bi-directional manner ^2, 5, 6^. CAFs are key effector cells in stroma, which produce ECM molecules such as collagen, fibronectin, laminin and secrete various cytokines and growth factors, which altogether stimulate tumor growth, angiogenesis, invasion, and metastases ^7, 8, 9^. CAFs mainly originate from pancreatic stellate cells (PSCs), the resident mesenchymal cells in pancreas ^10^. PSCs are normally present in low numbers in a quiescent form storing vitamin A droplets. However, during malignant transformation PSCs get activated and transform into myofibroblasts identified by α-smooth muscle actin (α-SMA) expression ^5^. Furthermore, PSCs have been shown to secrete growth factors that induce tumor cell growth, survival and migration ^11, 12^. In a recent study, we have also shown that TGFβ-activated PSCs stimulated tumor cell proliferation and endothelial cell activation, while inhibition of miR-199a/-214 in PSCs inhibited their pro-tumorigenic functions ^13^. Similarly, other studies have shown reprograming of the activated myofibroblasts in desmoplasia to normal state yields a better therapeutic response ^11^. By contrast, Özdemir *et al.* have demonstrated depleting αSMA^+^ myofibroblasts in PDAC genetic models resulted in aggressive tumors by compromising the immune system ^14^. In another study, Rhim *et al.* targeted deletion of sonic hedgehog (Shh) in fibroblasts led to aggressive tumor growth with undifferentiated phenotype ^15^. Although these studies suggest to take caution when targeting CAFs, new targets are desperately needed to design therapies to re-program CAFs to gain benefit for cancer therapy.

Integrins are heterodimeric transmembrane receptors consisting of α and β subunits, which form a large family of about 24 αβ integrins ^16, 17^. As cell adhesion receptors, integrins mediate cell-cell and cell-ECM interactions, but they also play an active role in signal transduction by regulating cytoskeletal organization, cell migration, proliferation and survival ^17, 18, 19^. Several integrins, αvβ3, αvβ5, αvβ6, α6β4, α4β1, α11β1 and α5β1 are reported to be overexpressed in various cancer types, being involved in tumor progression through tumor cell invasion and metastases ^20, 21, 22, 23, 24, 25^. The integrin α5 chain (ITGA5) forms a heterodimer with β1 and together they act as a receptor for fibronectin ^16, 22^. However, most studies have investigated integrins solely in relation to malignant tumor cells or vasculature^21, 22^ and there is scarce information on the significance of integrins in CAFs ^23^. However, significance of integrins and in particular ITGA5 in tumor stroma and CAFs is not well studied yet.

In this study, we investigated the prognostic and therapeutic role of ITGA5 in stroma of pancreatic tumors. We first evaluated the expression levels of ITGA5 in PDAC samples from patients and correlated them with the overall survival rate. Then, we examined the expression of ITGA5 in activated human PSCs *in vitro* and studied the impact of ITGA5 knockdown on the activation of PSCs and their phenotype. Next, we investigated the effect of ITGA5 knockdown in PSC-mediated tumor growth in a co-injection (PANC-1 + PSCs) tumor mouse xenograft model *in vivo*. Furthermore, in view of future clinical application, we designed a novel short peptidomimetic (AV3) against ITGA5 and examined its therapeutic efficacy *in vitro* and *in vivo* in the co-injection model and in patient-derived xenograft (PDX) models. Our study reveals ITGA5 as a key prognostic marker and a therapeutic target in pancreatic tumor. Importantly, our novel AV3 peptidomimetic inhibited the TGFβ-mediated ITGA5/FAK pathway *in vitro*. *In vivo*, AV3 reduced fibrosis and enhanced the efficacy of gemcitabine in PDX tumor model.

## Results

### Expression of ITGA5 in pancreatic tumor tissues and human pancreatic stellate cells

The immunohistochemical analyses for ITGA5 expression showed a strong expression in human PDAC samples, while there was no expression in normal pancreatic tissues (**Fig. 1A**). Within the tumor region, ITGA5 was largely and differentially expressed in the tumor stroma compared with tumor epithelial cells in which only a weak to negligible expression levels were observed (**Fig. 1A**). In the total sample cohort, ITGA5 was expressed in 66% of the PDAC patients, whereas α-SMA in 85% (**Table S1**). Since different populations of CAFs have been proposed in tumor stroma^26, 27^, we tried to identify varying CAF populations based on α-SMA and ITGA5 immunostainings. Interestingly, we identified three different populations: CAF1: α-SMA+/ITGA5+ (**Fig. 1B**, yellow colored cells), CAF2: α-SMA-/ITGA5+ (**Fig. 1B**, white arrows indicating red colored cells), and CAF3: α-SMA+/ITGA5- (**Fig. 1B**, green arrows indicating green colored cells). Out the total α-SMA+ CAF population, about 72% were positive for ITGA5 while 28% did not express ITGA5, indicating that there is a specific CAF population expressing ITGA5. Furthermore, both α-SMA and ITGA5 expression were associated (p=0.005; p=0.035, respectively) with the vascular invasion of the tumor. In univariate analyses, age, and sex did not demonstrate predictive values for overall survival (OS). However, pT-stage, pN-stage, margin status and ITGA5 expression in stromal cells were significantly predictive for OS. In multivariate analysis, only ITGA5 expression in stromal cells was significant prognostic factor for OS in pancreatic cancer (**Table 1**). Survival analysis reveals that the overexpression of both α-SMA and ITGA5 (log-rank p=0.022 and 0.008, respectively) were linked to significantly decreased overall survival (**Fig. 1C**). In addition, we examined the ITGA5 mRNA expression from public available dataset and found that ITGA5 expression in tumor was significantly higher than adjacent non-tumor tissue (**Fig. 1D**).

**Fig. 1.**
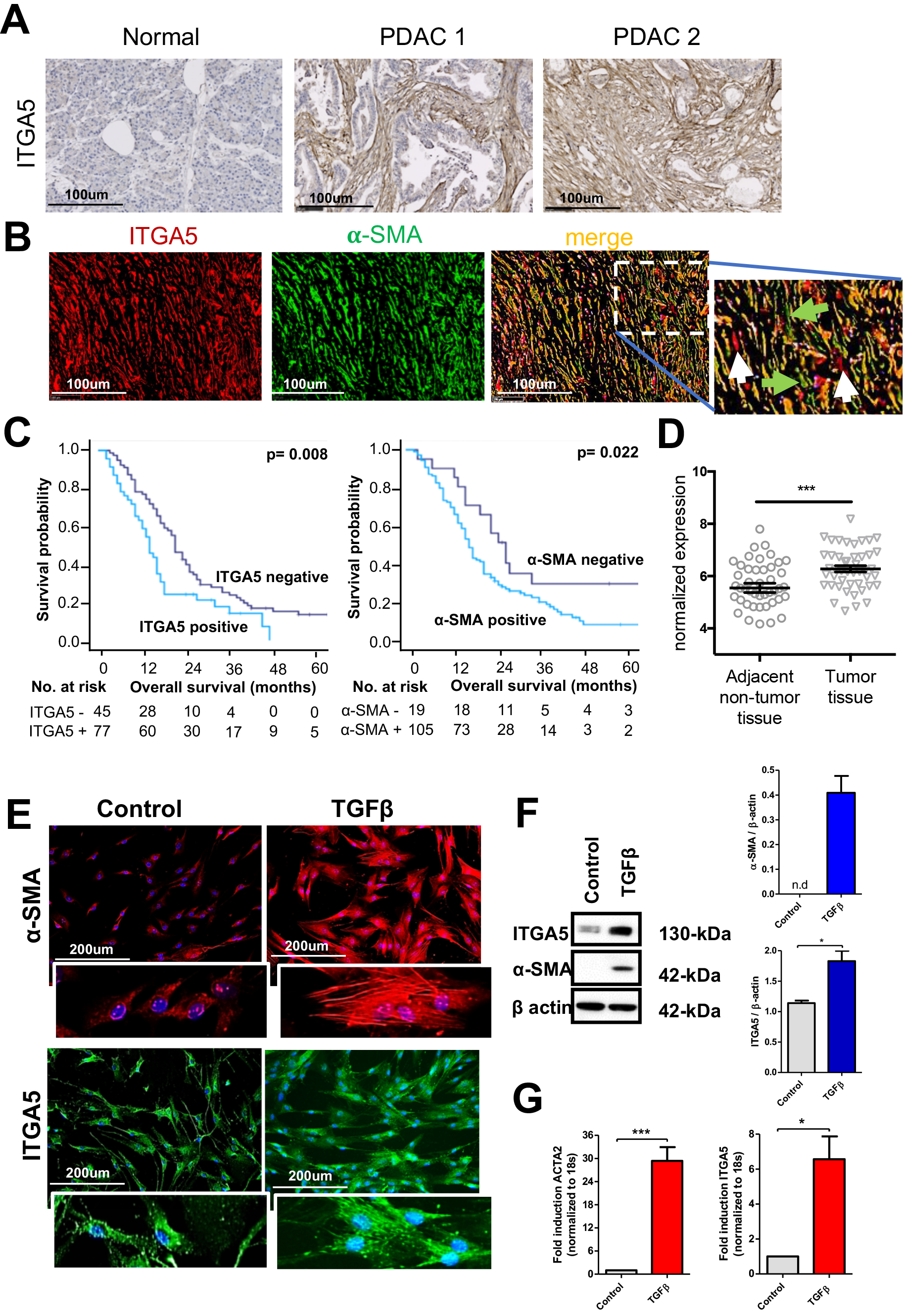
ITGA5 expression in pancreatic cancer. (**A**) Immunohistochemical (IHC) staining for ITGA5 performed on pancreatic tumor tissue microarrays and normal pancreas. Scale bar, 100μm. (**B**) Double immunofluorescent staining show ITGA5 (red) and α-SMA (green) with DAPI (blue nuclei) in the PDAC tissue. Scale bar, 100μm. (**C**) Kaplan-Meier overall survival (OS) curve for the stromal expression of ITGA5 and α-SMA in patients with PDAC. Log rank test was performed to calculate significant differences. Survival analyses was performed using the Kaplan-Meier method. (**D**) transcriptomic analysis of ITGA5 in public-available microarray dataset (GSE28735). (**E**) Immunofluorescent staining showing α-SMA and ITGA5 expression levels in hPSCs with or without TGFβ activation highlighting morphological changes. Scale bar, 200μm. (**F**) Western blot analysis and (**G**) gene expression analysis using qPCR for α-SMA and ITGA5 in hPSCs with or without TGFβ activation. Data represents mean ± s.e.m from at least three independent experiments; *p<0.05, ***p<0.001.

**Table 1.** Uni- and multivariable Cox proportional hazard model predictive value of ITGA5 expression on overall survival of patients with PDAC.

|  | Univariate |  |  | Multivariate |  |  |
| --- | --- | --- | --- | --- | --- | --- |
|  | HR | 95% CI | *p* | HR | 95% CI | *p* |
| Age ($\geq 65$ vs. <65 years) | 1.146 | 0.797-1.646 | 0.462 | | | |
| Sex (male vs. female) | 1.267 | 0.882-1.821 | 0.200 |  |  |  |
| pT-stage (pT2-4 vs. pT1) | 1.825 | 1.071-3.109 | **0.027** | 1.568 | 0.889-2.765 | 0.120 |
| pN-stage (pN1 vs. pN0) | 1.557 | 1.008-2.404 | **0.046** | 1.228 | 0.768-1.962 | 0.391 |
| Margin status (R1 vs. R0) | 1.681 | 1.133-2.495 | **0.010** | 1.349 | 0.874-2.080 | 0.176 |
| ITGA5 (High vs. low expression) | 1.870 | 1.267-2.762 | **0.002** | 1.591 | 1.056-2.399 | **0.026** |

Abbreviations: CI, Confidence interval; HR, hazard ratio; ITGA5, Integrin alpha 5; Significant P values are bold.

PSCs are considered as the main source for CAFs in pancreatic tumor stroma ^6^. Upon activation, PSCs differentiate into α-SMA expressing myofibroblast-like cells and produce abundant ECM^4^. In this study, we show that quiescent human PSCs (hPSCs) were α-SMA negative and after activation with human recombinant TGFβ, they developed actin filaments fibers and showed high expression levels of α-SMA, as shown with immunofluorescence staining (**Fig. 1E**). These data were also confirmed at protein and mRNA levels using western blot and qPCR analyses (**Fig. 1F, 1G**). However, α-SMA is barely detectable in quiescent hPSCs (**Fig. 1F**). Furthermore, we found that, in contrast of α-SMA, ITGA5 was also expressed by quiescent hPSCs and its expression was significantly upregulated after TGFβ activation, as shown with immunofluorescent staining, western blot and qPCR analyses (**Fig. 1, E-G**). Importantly, expression of ITGA5 along the actin filaments indicates towards its potential involvement in the attachment and maintenance of morphology of the activated hPSCs. To this end, TGFβ-activated hPSCs seem to be like CAF1 (α-SMA+/ITGA5+), while quiescent hPSCs at least showed some expression levels of CAF2 (α-SMA-/ITGA5+). Yet, CAF3 (α-SMA+/ITGA5-) population is not understood here which might result from an alternative activation.

### ITGA5 knockdown attenuates TGFβ-induced PSC activation and differentiation

To study the effect of ITGA5 on the activation of hPSCs, we knocked down its expression using puromycin-resistant lentiviral shRNA plasmid. The stably ITGA5-knockdown (sh-ITGA5) hPSCs showed a reduced ITGA5 and α-SMA expression levels compared to the negative control (NC) shRNA (sh-NC) (**Fig. 2, A-C**). As shown in **Fig. 2A** (zoomed images), the overexpression of ITGA5 along the actin filaments in the TGF-β-activated hPSCs was lost in sh-ITGA5 hPSCs. Moreover, the phalloidin staining suggests sh-ITGA5 hPSCs turned into large flat cells compared to sh-NC stretched cells, which is likely due to the loss of cytoskeletal filaments (**Fig. 2D**). We also found that there was a significant reduction in TGF-β1-induced collagen-1 expression in sh-ITGA5 hPSCs (**Fig. S1A**). Furthermore, sh-ITGA5 hPSCs had a significantly reduced mRNA expression of key activation markers such as ACTA2 (α-SMA), Fibronectin1 (FN1), Periostin (POSTN) (**Fig. 2E**), Platelet-derived growth factor beta receptor (PDGFβR) (**Fig. S1B**). Since the activated hPSCs produce abundant ECM during tumorigenesis and attach via adhesion proteins, we investigated the impact of ITGA5 knockdown on ECM and adhesion proteins using a human profiler gene array comprised of 84 genes. As shown in the **Fig. 2F** and **Table S4**, the expression levels of several genes related to ECM and adhesion molecules were significantly downregulated (fold>0.5) in sh-ITGA5 compared to sh-NC hPSCs. Activation of sh-NC hPSCs with TGF-β led to the upregulation (fold >2.0) of about 42 genes.

**Fig. 2.**
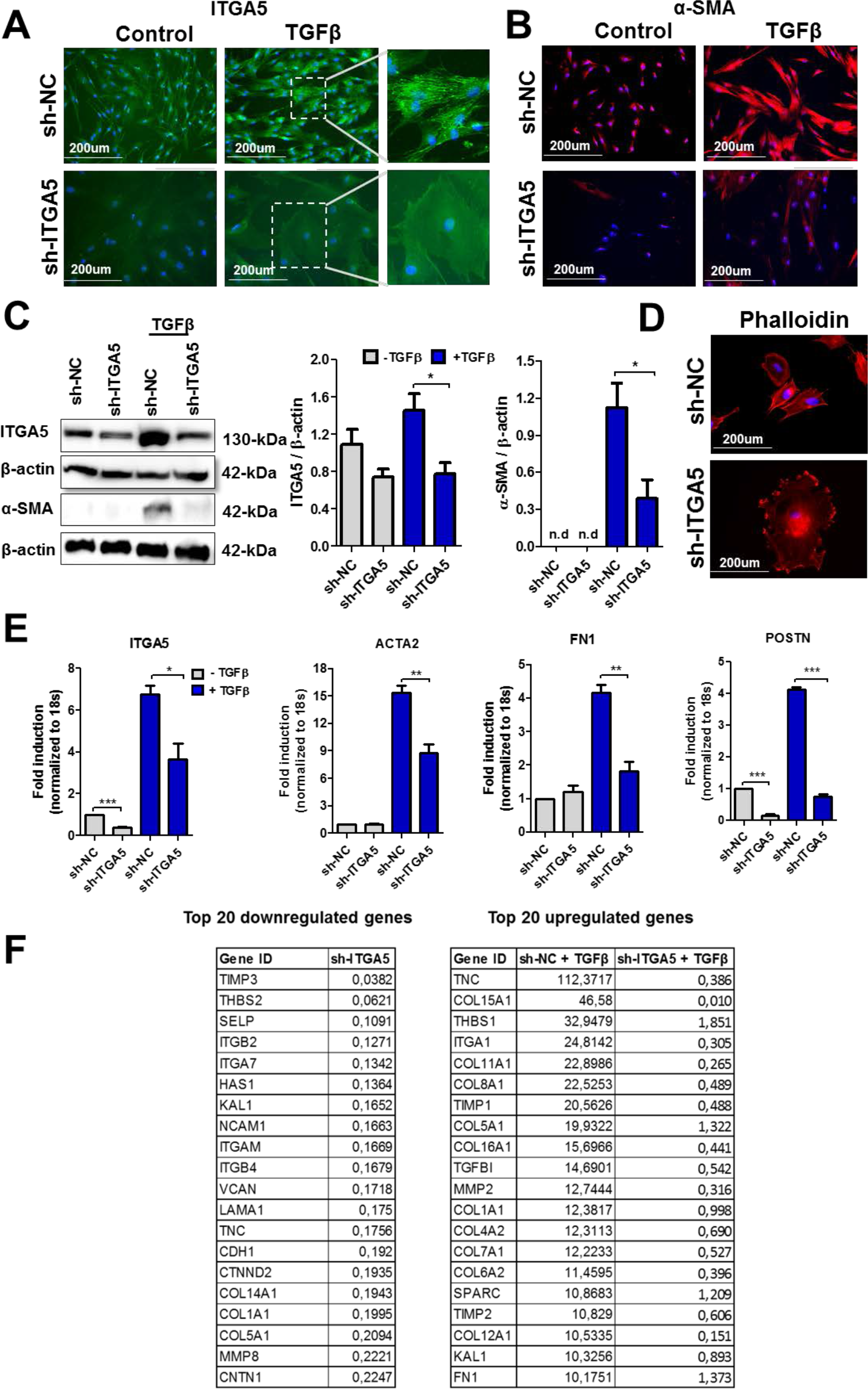
ITGA5 knockdown attenuates hPSCs activation and differentiation. (**A-C**) Immunofluorescent staining and western blot analyses show high or low ITGA5 and α-SMA expression in hPSCs after ITGA5 knockdown (sh-ITGA5) compared to negative control (sh-NC) hPSCs with or without TGFβ activation. (**D**) sh-ITGA5 hPSCs show morphological differences compared to sh-NC hPSCs. (**E**) The expression levels of ITGA5, ACTA2, Fibronectin1, Periostin were determined by qPCR. The expression levels of each gene were normalised by that of 18S. Data represents mean ± s.e.m from at least three independent experiments; *p<0.05, **p<0.01, ***p<0.001. (**F**) The differential gene expression of adhesion molecules and ECM using human ECM RT^2^ Profiler™ PCR Array in sh-ITGA5 with or without TGFβ activation. In the left table, sh-ITGA5 data is relative to sh-NC (set at 1.0) showing top 20 downregulated genes after ITGA5 knockdown. In the right table, sh-NC + TGFβ data are relative to sh-NC hPSCs while sh-ITGA5 + TGFβ data are relative to sh-ITGA5 hPSCs, showing the most induced genes after TGFβ activation which are not induced after ITGA5 knockdown.

Interestingly, activation of sh-ITGA5 hPSCs with TGFβ did not upregulate the downregulated genes (**Fig.2F** and **Data file S1**). These data indicate that the downregulation of ITGA5 expression level leads to inhibition of TGFβ-mediated differentiation of hPSCs into myofibroblasts-like cells and thereby inhibit ECM production.

### ITGA5 knockdown inhibits hPSC phenotype and activation via TGFβ-induced FAK/Smad/AKT pathways

Integrins are known to control various cellular processes such as migration, cell adhesion, contraction and proliferation ^17^. We therefore investigated whether ITGA5 plays a role in controlling PSC phenotype as shown in **Fig. 3**. We examined the cell-to-ECM and cell-to-cell adhesion using a plate cell-adhesion assay and a spheroid assay, respectively. Since ITGA5 is a receptor for fibronectin, we examined the attachment of cells on fibronectin-coated plates. Compared with sh-NC hPSCs, the sh-ITGA5 hPSCs attached significantly lesser to fibronectin (**Fig. 3A**) and also formed significantly less compact spheroids after 6 days of culturing (**Fig. 3B**). The effect of ITGA5 knockdown on cell proliferation and TGFβ-induced contractility was investigated using BrdU assay and 3D collagen gel contraction assay. sh-ITGA5 hPSCs showed a reduction in proliferation and contractility (**Fig. 3C and 3D)**. In addition, we examined the effect of ITGA5 knockdown on migration of hPSCs using a scratch assay and found a significantly lower migration rate (60%, p<0.01) compared with sh-NC hPSCs (**Fig. 3E**). This reduction in migration was not related to the effect on proliferation, as this assay was performed within 15h while there was no change in proliferation up to 24h **(Fig. 3C)**. These data suggest that ITGA5 plays a crucial role in regulation of cell adhesion, migration, proliferation, and TGFβ-induced contractility of hPSCs. Having established that ITGA5 knockdown attenuates TGFβ induced differentiation and phenotypic changes in hPSCs, we examined the potential pathways responsible for these activities. We investigated the effect on the TGFβ signaling pathways i.e. pSmad2 and pAKT. In addition, ITGA5 is known to act via Focal Adhesion Kinase (FAK) as a canonical pathway and its crosstalk with TGFβ has also been proposed in literature ^28, 29^. We therefore also investigated the effect of ITGA5 knockdown on FAK pathway. We performed western blots at early (30 min) and late (48h) time points after TGFβ activation. We found that pSmad2 expression was induced at 30 min after TGFβ activation (**Fig. 3F**), while pAKT (phosphorylation at S473 site) and pFAK (phosphorylation at Y397 site) were induced only after 48h (**Fig. 3G**). Interestingly, ITGA5 knockdown did not only reduce the phosphorylation of Smad2 (**Fig. 3F**), but also AKT and FAK pathways in both non-activated and TGFβ activated hPSCs (**Fig. 3F**).

**Fig. 3.**
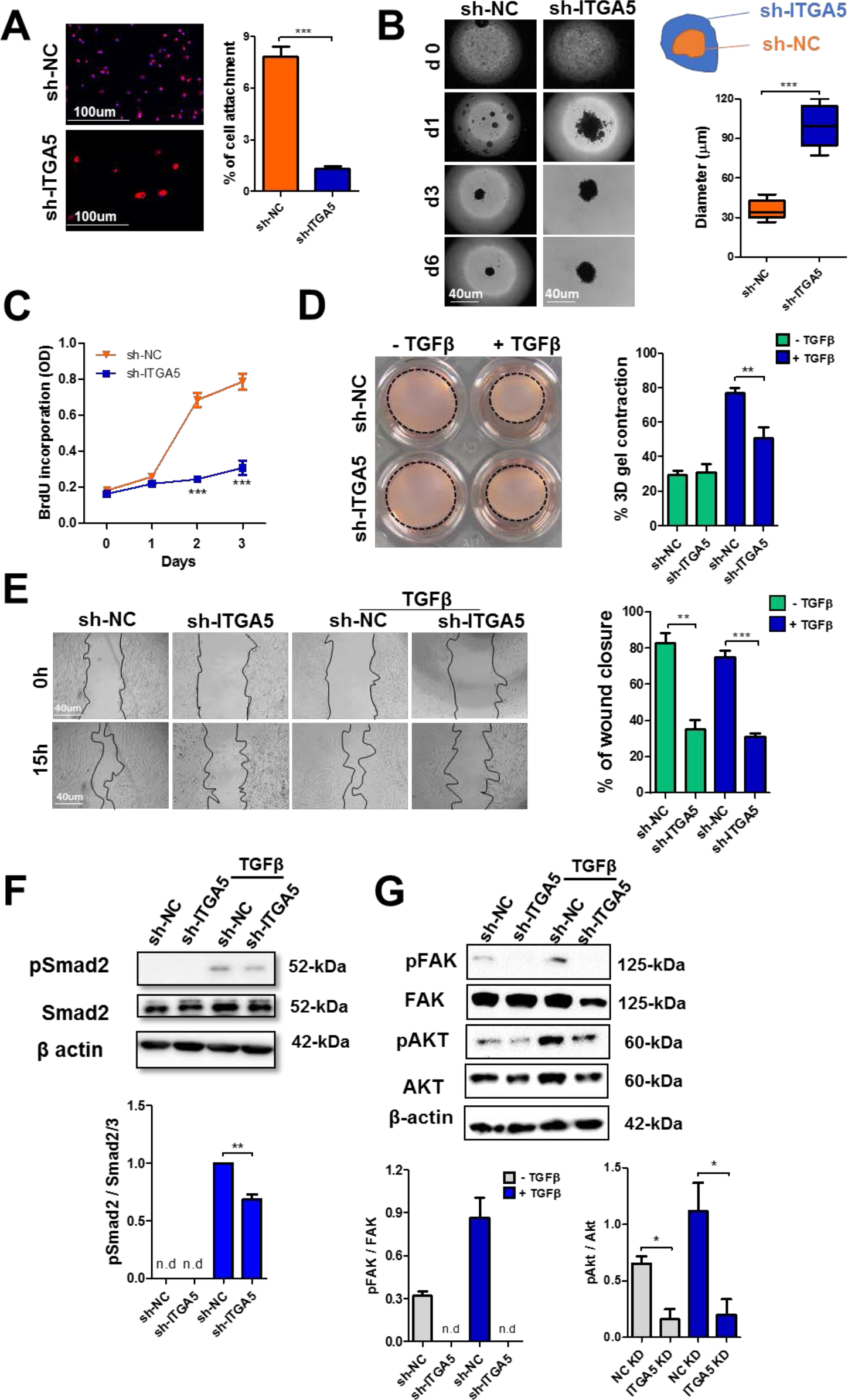
ITGA5 knockdown inhibits hPSCs activation. (**A**) Cell adhesion assay was performed on fibronectin coated plate, sh-ITGA5 hPSCs adhere less to fibronectin in comparison to sh-NC cells. (**B**) Spheroids are prepared either from sh-ITGA5 or sh-NC hPSCs using hanging drop method. (**C**) Cell proliferation was assessed by BrdU incorporation ELISA, and the absorbance at 370nm of the samples is shown at different days. (**D**) Representative images from 3D collagen gel contractility assay show less TGFβ induced contractility in sh-ITGA5 hPSCs after 96 h. (**E**) Representative microscopic images of cell migration after 15 h and quantitative analyses show sh-ITGA5 hPSCs migrate less compared to that of sh-NC with or without TGFβ activation. Western blot analyses show the expression levels of p-Smad2/Smad2 at t=30 min (**F**) and of pFAK-Y397/FAK and p-AKT/AKT at t=48h (**G**) after TGFβ incubation in sh-ITGA5 and sh-NC hPSCs. Densitometry analyses were performed using Image J software. Data represents mean ± s.e.m from at least three independent experiments; *p<0.05, **p<0.01, ***p<0.001.

Both pAKT and pFAK were not activated by TGFβ at t=30 min (data not shown). In addition, we examined the effect of ITGA5 knockdown on ITGA5-induced direct signaling molecules i.e. FHL3 (four and a half LIM domain protein 3) and Paxillin and downstream genes i.e. Rho family proteins Cdc42, Rac1 and RhoA. Importantly, the expression levels of these genes, either without or with TGFβ activation, were significantly reduced in ITGA5 knockdown hPSCs (**Fig. S1B**). These findings demonstrate that ITGA5 controls the activation of PSCs via both canonical (FAK) and non-canonical TGFβ-induced signaling pathways.

### ITGA5 knockdown inhibits PSC-induced paracrine effect *in vitro* and tumor growth *in vivo*

Differentiated PSCs or CAFs secrete growth factors and cytokines which stimulate tumor cells for their proliferation and migration, as depicted in **Fig. 4A**. To mimic this process, we collected conditioned media from sh-ITGA5 and sh-NC hPSCs with or without TGFβ activation. In our previous study, we have shown PANC1 tumor cells treated with conditioned media collected from TGFβ-activated hPSCs displayed higher tumor cell growth^13^. In this study, we found the similar paracrine effect of the TGFβ-activated hPSCs on PANC-1 cells (**Fig. 4B**). Interestingly, conditioned media obtained from TGFβ-activated sh-ITGA5 hPSCs did not induce the growth of tumor cells (**Fig. 4B**). Likewise, we evaluated the effect of sh-ITGA5 and sh-NC hPSCs conditioned media on the migration of PANC-1 tumor cells using a transwell migration assay. Notably, tumor cells with sh-ITGA5 hPSCs conditioned media clearly showed lower migration than those treated with the sh-NC hPSCs conditioned media (**Fig. 4C**). In relation to that, we found that knockdown of ITGA5 led to reduction of CXCL12, IL-6 and TGFβ genes in non-activated/activated hPSCs (**Fig. S1B**), explaining the inhibition of paracrine effects by sh-ITGA5 hPSCs. These data are clearly suggesting that knockdown of ITGA5 in hPSCs inhibits the paracrine interactions between PSCs and pancreatic cancer cells.

**Fig. 4.**
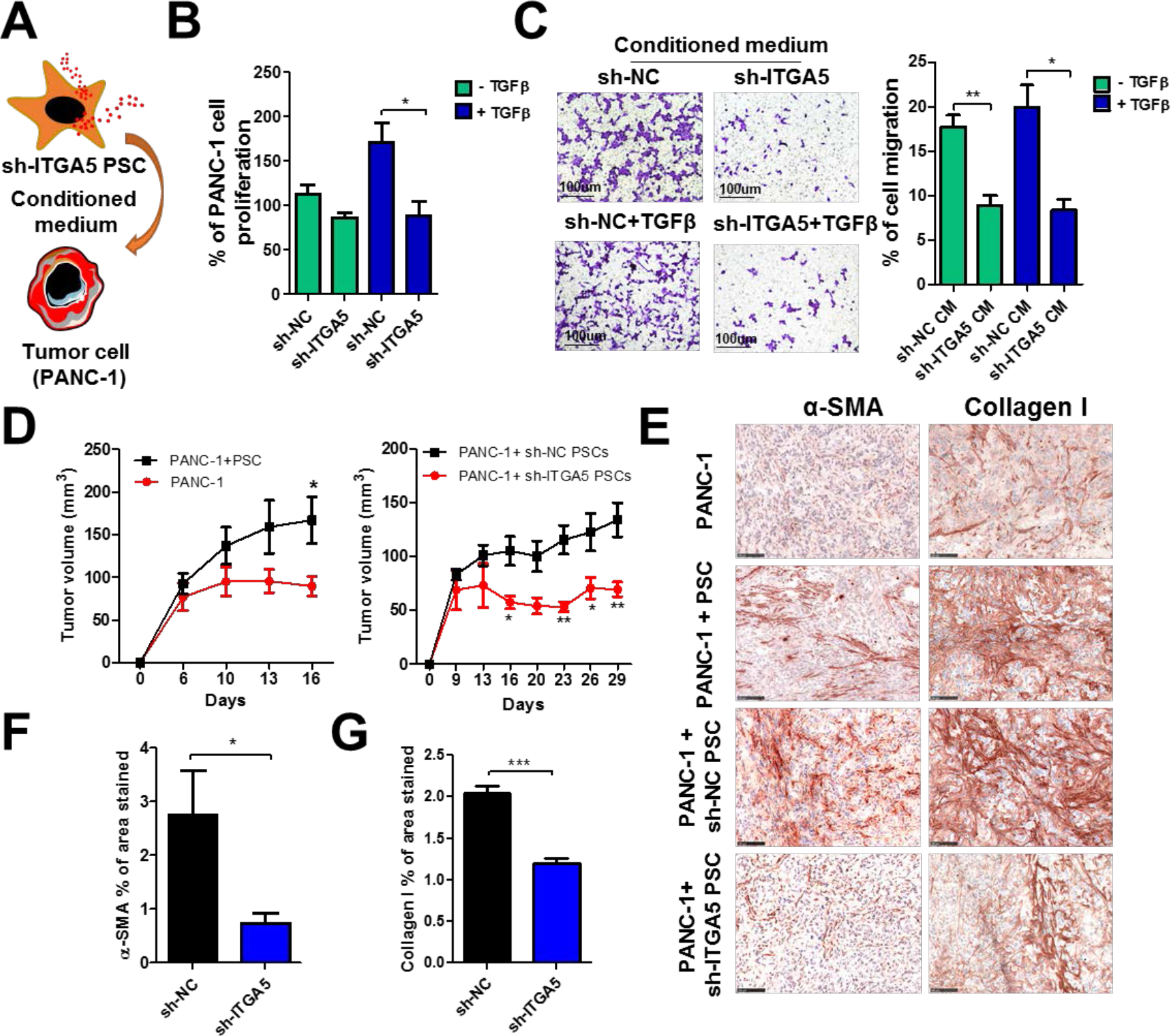
ITGA5 knockdown inhibits PSC-induced paracrine effects. (**A**) Schematic representation of paracrine effect of PSCs on tumor cells. (**B**) PANC-1 tumor cell growth was assessed by Alamar blue after incubating with conditioned medium obtained either from with or without TGFβ activated sh-ITGA5, sh-NC hPSCs. (**C**) Representative images from transwell migration assay show conditioned medium (CM) from sh-ITGA5 hPSCs either with or without TGFβ shows less migration on PANC-1 compared to that of sh-NC CM either with or without TGFβ. (**D**) Tumor growth curves from a co-injection of the PANC-1 versus PANC-1 + hPSCs (left graph) and Panc-1 + sh-NC hPSCs versus PANC-1 + sh-ITGA5 hPSCs. (**E**) Microscopic pictures show immunohistochemical stainings for α-SMA and Collagen-I in tumors, and (**F**) and (**G**) show the quantitation of the α-SMA and Collagen-I stainings, respectively. Data represents mean ± s.e.m; *p<0.05, **p<0.01, ***p<0.001.

To investigate whether knockdown of ITGA5 in hPSCs retards their pro-tumorigenic effects *in vivo*, we established a co-injection tumor model in immunodeficient SCID mice. As consistent with previous reports ^12, 30^, tumors derived from co-injection of PANC-1 and hPSCs showed a significant increased tumor growth compared to tumors with PANC-1 cells alone (**Fig. 4D**). Furthermore, immunohistological examination revealed that PANC-1+hPSCs tumors were highly fibrotic, as indicated by overexpression of α-SMA and collagen I, in comparison with PANC-1 tumors (**Fig. 4E**). In addition, we found that tumors derived from co-injection had higher expression levels of ITGA5 compared to PANC-1 tumors (data not shown). Subsequently, we induced tumors by co-injecting PANC-1 with hPSCs, either stably transfected with sh-NC or sh-ITGA5 lentiviral plasmids. The sh-ITGA5 hPSCs had a stable knockdown of 55% (data not shown). As shown in **Fig. 4D**, we found that PANC-1+ hPSCs (sh-ITGA5) tumors had a significantly slower tumor growth compared to that of PANC-1+ hPSCs (sh-NC) (**Fig. 4D**). Furthermore, tumors with ITGA5-knockdown hPSCs showed significantly less fibrosis area compared to tumors with normal hPSCs, as shown with immunohistochemical staining of α-SMA and collagen I (**Fig. 4, E-G**). These findings suggest that ITGA5 in hPSCs play a key role in inducing its pro-tumorigenic actions *in vivo*.

### Novel ITGA5 blocking peptidomimetic AV3 inhibits hPSC differentiation

To develop a therapeutic peptidomimetic specifically against ITGA5, overlapping sequences (12 amino acids (a.a.) long with 8 a.a. overlaps) from human FN-III domains-9, and 10 were synthesized and displayed on a cellular membrane. Domains 9 and 10 of FN-III, are reported to be responsible for binding to the α5β1 receptor, as shown with the docking experiments elsewhere^31^. With the interaction studies between the identified sequences and α5β1 receptor, we found two consecutive sequences “TTVRYYRITYGE” and “YYRITYGETGGN” strongly bound to the receptor but the sequences immediately before or after these sequences did not show any binding. From these analyses, we concluded the common sequence RYYRITY (named here as AV3) as the minimal sequence (**Fig. 5A)**, responsible for the binding to α5β1. The binding studies with peptidomimetic AV3 conjugated with 5-FAM fluorescent dye via a PEG linker (AV3-PEG(6)-FAM) showed a strong binding to hPSCs with increasing concentrations. The binding was enhanced in TGF-β-activated hPSCs, as shown with flow cytometry as well as by fluorescent microscopy (**Fig. 5B and 5C**). Surprisingly, we found no difference in binding at lower concentrations. It is likely that α5β1 receptor was sufficiently expressed in both non-activated and activated hPSCs to accommodate low concentrations of AV3-PEG(6)-FAM equally.

**Fig. 5.**
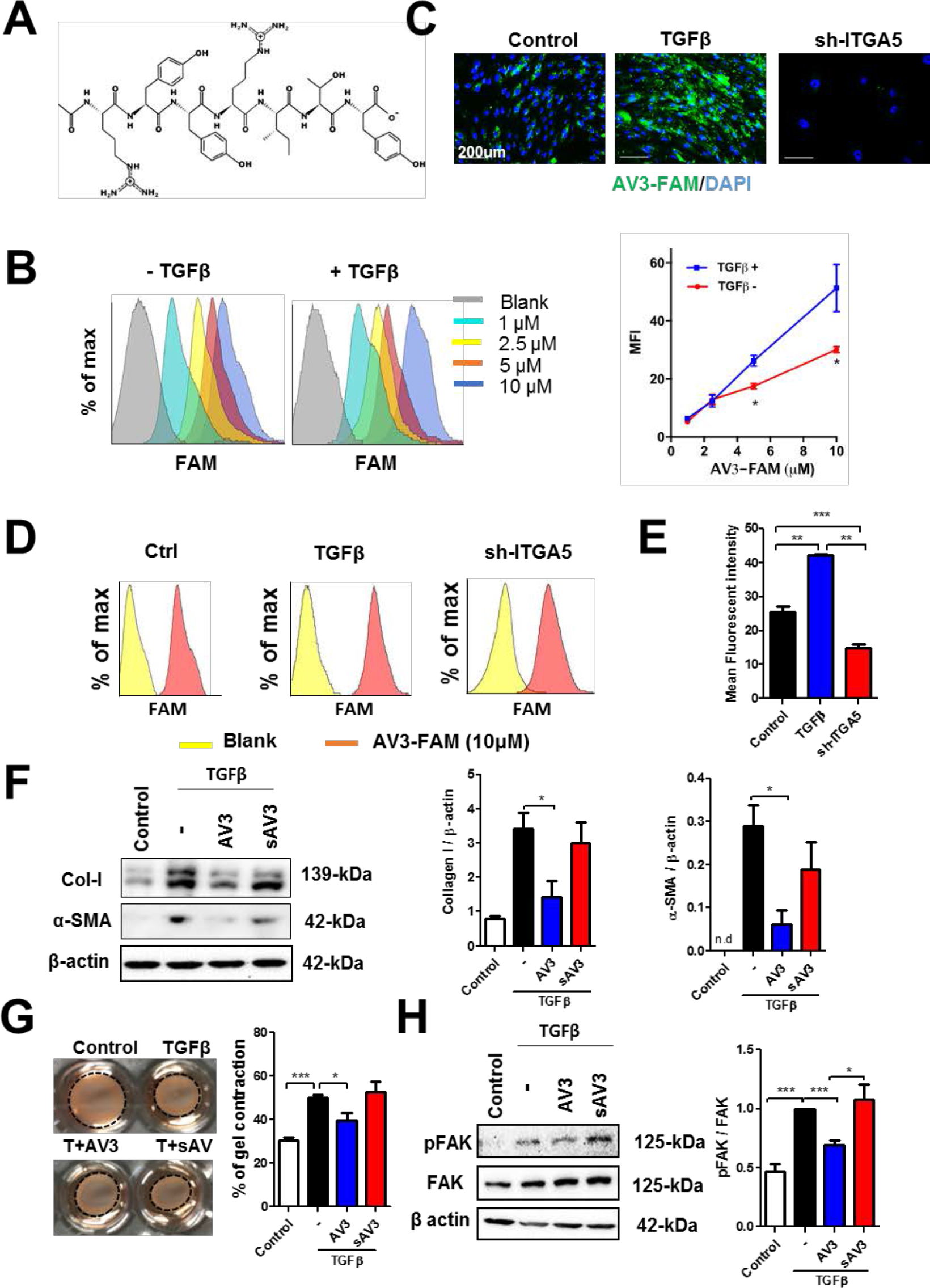
A Novel peptidomimetic against ITGA5. (**A**) Chemical structure of AV3 peptidomimetic. (**B**) Flow cytometry histograms and analysis show an increased binding of AV3-PEG(5)-FAM with the increasing concentration in TGFβ activated hPSCs compared to that of control (non-activated) hPSCs. Binding of AV3-FAM in control hPSCs, TGFβ-activated hPSCs and sh-ITGA5 hPSCs as shown in the representative microscopic fluorescent images (**C**), flow cytometry histograms (**D**) and their quantitative analysis (**E**). (**F**) Western blot analyses showing that AV3 inhibits α-SMA and collagen 1 expression levels in hPSCs, whereas scrambled (s) AV3 shows no inhibitory effects (**G**) Representative images from collagen gel assay show AV3 peptidomimetic inhibits TGFβ induced collagen gel contractility after 72 h. (**H**) Western blot analyses show the expression levels of pFAK, FAK and β-actin in hPSCs following AV3 treatment in 8 hr TGFβ activated hPSCs. Data represents mean ± s.e.m from at least three independent experiments; *p<0.05, **p<0.01, ***p<0.001.

In contrast, at higher concentrations the differences became apparent. The increase in binding was in line with the increased ITGA5 expression after TGFβ activation (**Fig. 1D and 1E**). To study the specificity for ITGA5, we examined the binding to sh-ITGA5 hPSCs and found a substantial loss of binding (**Fig. 5, C-E**). Importantly, the loss of binding was in line with the inhibition of ITGA5 in knockdown cells (**Fig. 2C**). These data confirm the selection of a short endogenous peptidomimetic sequence specifically binding ITGA5.

Furthermore, we investigated whether blocking ITGA5 with AV3 could inhibit TGFβ-mediated activation of hPSCs. Interestingly, AV3 significantly reduced the expression levels of differentiation marker ACTA2 (α-SMA) and Collagen I both at the transcription and translational level using western blot analysis and immunocytochemical staining (**Fig. 5F, S2A and S2B**). We also examined whether AV3 could inactivate already activated CAFs isolated from patients. We observed a clear inactivation of the primary pancreatic CAFs, as shown with immunocytochemical stainings for α-SMA and collagen I (**Fig. S2C**). In contrast, scrambled AV3 did not show any inhibitory effects. Furthermore, we examined the effect of the AV3 on the TGFβ-induced contractility of hPSCs in 3D collagen gel. In line with the ITGA5 knockdown data, we found that AV3 peptidomimetic significantly inhibited the TGFβ-induced contractility of hPSCs after 72h, as shown in **Fig. 5G**.

Since AV3 inhibited the TGFβ-mediated effects, we were interested in understanding the mechanism of action of AV3. In Fig.3f, we showed that TGF-β activated the integrin signaling i.e. phosphorylation of FAK and knockdown of ITGA5 inhibited this signaling. We seek whether interaction of AV3 with ITGA5 could inhibit the TGFβ-mediated direct activation of ITGA5 signaling via the FAK pathway. We found that within 8h of incubation with TGFβ, the FAK pathway was activated in hPSCs and interestingly, treatment with AV3 significantly inhibited this activation (**Fig. 5H**). This is in line of the sh-ITGA5 data (**Fig. 3F**). However, in contrast, we observed no effect on pSmad2 with AV3 treatment (**Fig. S2D**), which clearly distinguishes between the receptor deletion and the receptor blocking techniques.

### *In vivo* AV3 potentiates the effect of gemcitabine *in vivo*

As shown in **Fig. 4D**, we already demonstrated that co-injection of hPSCs with PANC-1 tumor cells stimulated tumor growth by inducing fibrosis and that knockdown of ITGA5 in hPSCs restricted this enhancement of the tumor growth. We therefore investigated whether AV3 peptidomimetic was able to inhibit the hPSC-induced tumor growth in this model. Remarkably, treatment with AV3 reduced the tumor growth significantly after either intraperitoneal (i.p) or local intratumoral (i.t) injection (**Fig. S3**). In contrast, the scrambled peptidomimetic (sAV3) did not show any inhibitory effect. We found that the treated tumors had reduced fibrosis, as shown with immunostainings for collagen I (**Fig. S3B**) and fibroblast activation markers such as desmin (**Fig. S3C,D**). We examined the tumor vasculature using the endothelial cell marker CD31 but found no effect of the treatment on the tumor vasculature (**Fig. S3E**). Furthermore, AV3 was well tolerated by the animals, as there was no change in the body weight after multiple treatments (**Fig. S5A and S5B**).

Having seen the anti-fibrotic effect of AV3 *in vivo*, we extended our investigation in a patient-derived xenograft (PDX) pancreatic tumor model. As shown in **Fig. 6A**, a small piece of the pancreatic tumor was isolated from a patient and grown to passage 2 (P2) followed by implanting into the flank of a NOD/SCID mice. To confirm that the used tumor had abundant stroma and expressed ITGA5 and, we examined the P0 tumor (tumor from the patient-193) for the expression of ITGA5, α-SMA and collagen-I immunohistochemically and found a high expression of these markers (**Fig. S4**). AV3 alone did not show reduction in tumor growth which might be due to replacement of patient tumor stroma by mouse fibroblasts, which likely have different phenotype and do not contribute to the tumor growth. As expected, treatment with gemcitabine reduced the growth of the PDX significantly, but co-treatment with AV3 and gemcitabine reduced the tumor growth even more (**Fig. 6B**). Furthermore, these results were confirmed by the weight of the isolated tumors at the end of the experiment (**Fig. 6C**). In this model, AV3 alone did not show significant tumor growth inhibition. The enhanced effect of gemcitabine was most likely due to its better perfusion and delivery. To confirm the effect of AV3 on the tumor perfusion, we performed tumor distribution of the indocyanine green (ICG) dye using optical imaging and found that indeed treatment with AV3 significantly enhanced the tumor perfusion, as can be seen with higher accumulation of ICG dye in AV3-treated tumors (**Fig. 6D**). Furthermore, we examined the isolated tumors to see the effect on fibrosis and found that the treatment with AV3 alone or combined treatment reduced the expression of α-SMA and Collagen I (**Fig. 6E**).

**Fig. 6.**
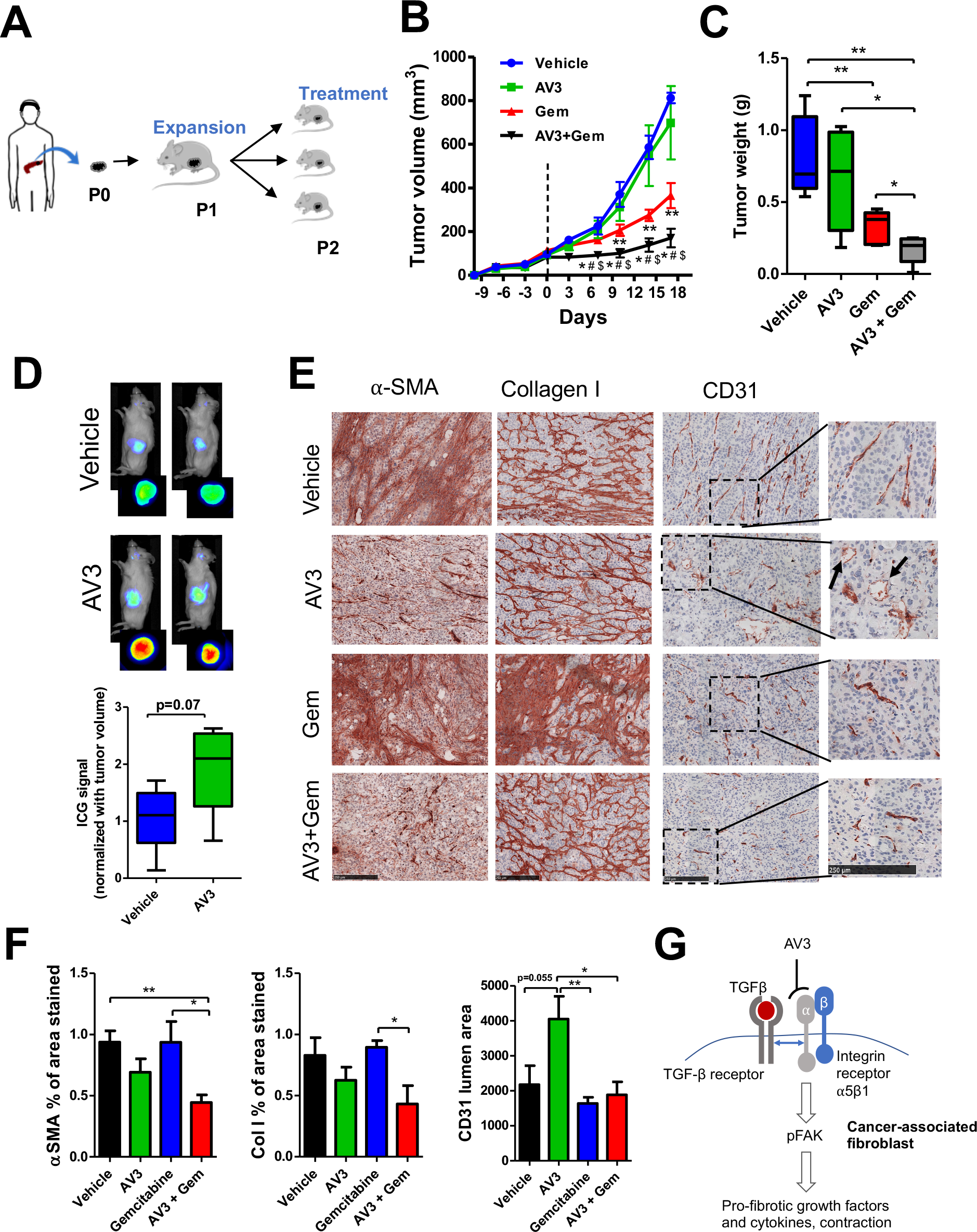
AV3 potentiates the anti-tumor effect of gemcitabine in PDX model in mice. (**A**) Schematic representation of the generation of human pancreatic patient-derived xenograft (PDX) tumor model. (**B**) Tumor growth curves in PDX model after the treatment with vehicle, AV3 (20 mg/kg, i.p., twice a week), gemcitabine (Gem), or AV3 + Gem. The doses of AV3 and Gem was (20 mg/kg and 50 mg/kg, respectively) i.p., twice a week started at day 0 (when tumors reached ~150mm^3^). Data are represented as mean ± s.e.m. * p<0.05 Gem vs. AV3+Gem, # p<0.05 AV3 vs. AV3+Gem, $p<0.05 Vehicle vs. AV3+Gem. **p<0.05 Vehicle vs Gem (**C**) Isolated tumor organs weight at the end of the experiment after different treatments as described above. **(D)** Optical imaging showing the accumulation of ICG dye in tumors treated with either vehicle or AV3 peptidomimetic in PDX model. Mice were injected with ICG dye via tail vein at the dose of 5mg/kg. After 24 hr of the injection, tumors were imaged externally or after isolation using a small animal imager. The ICG fluorescence signal was quantified. n=5 mice per group. Data represent mean ± s.e.m. (**E**) Microscopic pictures of immunohistochemical staining α-SMA, collagen-I and CD31 and their quantitation (**F**) showing the effect of different treatments on tumors. Data represents mean ± s.e.m; *p<0.05, **p<0.01. (**G**) The schematic diagram depicting the mechanism of action of AV3 peptidomimetic. TGF-β activates TGF-βR, which interacts with ITGA5 receptor and initiates FAK signaling. AV3 binds to ITGA5 and blocks TGF-β-induced ITGA5/FAK signaling.

Of note, gemcitabine alone did not show these effects. Moreover, we found that the blood vessel lumens in AV3-treated tumors were decompressed and opened up, as shown in **Fig. 6E** (see arrows) and **6F**, suggesting that the reduction in fibrosis supported normalization of blood vessels and thereby induced delivery of gemcitabine and the anti-tumor effects. We also show that treatment with AV3 peptidomimetic did not show any toxic effects as can be seen with the body weight, liver and lungs weight but the mean liver and lung weights of the AV3 and gemcitabine combined group were equal to that of the normal mice (**Fig. S5C-E**).

## Discussion

The abundant desmoplastic stroma reaction in pancreatic tumors has been recognized for inducing aggressive tumor growth, distant metastasis, resistance to drug therapy and acting as a barrier to drug delivery ^32^. Targets within the tumor stroma are therefore under intensive investigation to improve prognosis by developing novel therapeutics to hamper stromal tumor-promoting function. In the present study, we highlight integrin α5 (ITGA5) as a key target in tumor stroma showing its significance in prognosis of pancreatic cancer. Our ITGA5 knockdown studies show that ITGA5 is crucial for the maintenance of the PSCs phenotype and controls TGFβ-mediated activation through canonical FAK pathway and non-canonical TGFβ signaling pathways including Smad2 and AKT. Furthermore, knockdown of ITGA5 inhibits PSC-induced tumor cell growth and migration *in vitro* and PSC-driven tumor growth *in vivo*. A novel ITGA5 binding peptidomimetic sequence (AV3) has been developed which show inhibition of TGFβ-mediated activation of hPSCs *in vitro*. *In vivo*, AV3 reduces fibrosis in tumor stroma established in xenograft as well as PDX tumor models. In the PDX model, addition of AV3 to gemcitabine-based chemotherapy enhanced the anti-tumor effects strongly by reducing the tumor stroma. Being ECM receptors, integrins are highly upregulated on the membrane of myofibroblasts, the main producers of ECM molecules ^17, 21, 28^. Targeting different integrins on myofibroblasts has been shown to inhibit fibrosis in multiple organs such as liver, lung and kidney ^33, 34^. ITGA5, as subunit of the fibronectin receptor, has been reported to be overexpressed in activated fibroblasts in fibrosis and cancer as well as in tumor epithelial cells undergoing epithelial-mesenchymal transition ^22, 35, 36^.

In this study, overexpression of ITGA5 was observed in stromal CAFs in clinical samples of pancreatic cancer, with a very weak expression in malignant cells, confirming the specificity of ITGA5 for CAFs. In literature, high expression levels of CAF markers such as α-SMA and PDGFβR in pancreatic tumor have been shown to be associated with poor prognosis ^37, 38^. In line with these data, we indicate ITGA5 as a novel stromal prognostic marker, as its high expression was independently associated with poor overall survival. In literature, different CAF populations have been proposed ^26^ and based on these data, we could identify three different populations of CAFs (CAF1: α-SMA+/ ITGA5+, CAF2: α-SMA-/ ITGA5+ and CAF3: α-SMA+/ ITGA5-). Yet, the largest population was of CAF1.

PSCs are regarded as the main source of CAFs ^11^ and upon activation with TGFβ they acquire myofibroblastic phenotype, as shown earlier by us and others ^13, 39, 40^. TGFβ-mediated activation of hPSCs *in vitro* resulted in stretched and elongated α-SMA expressing myofibroblasts which overexpressed ITGA5 compared to quiescent cells. Of note, quiescent hPSCs were α-SMA negative but yet strongly positive for ITGA5, resembling the features of the CAF2 population found in tumor material. These data indicate the potential role of ITGA5 in the maintenance of hPSCs phenotype. Knockdown of ITGA5 using shRNA, as shown in this study, led to the reprogramming of hPSCs inhibiting TGFβ-induced differentiation into myofibroblasts and ECM production. Notably, ITGA5 knockdown hPSCs became flattened due to the loss of stress filaments and had reduced cell-to-cell adhesion, migration, proliferation and contractile properties. These results can be explained as integrins are responsible for maintaining cell phenotype such as adhesion, migration, contraction and proliferation ^16^.

Besides the change in the phenotype, knockdown of ITGA5 also inhibited TGFβ-induced activation of hPSCs. In literature, integrins have been reported to interact with several growth factor receptors, including PDGFR-β, c-Met, VEGFR, EGFR, and TGFβ receptors ^41^. On the one hand, TGFβ1 is shown to induce the expression of α5β1 and activate FAK signaling due to ligation and clustering of integrins ^42^. On the other hand, α5β1 integrin is also shown to modulate TGFβ1/Smad pathways directly or indirectly via different mechanisms ^28, 29^. In line with these data, we herein show that TGFβ1 not only activates its own signaling pathways (i.e. pSmad2 and pAKT), but also the ITGA5-mediated pFAK signaling pathway. Interestingly, ITGA5 knockdown in hPSCs inhibited TGFβ-mediated both early (pSmad2) and late (pAKT and pFAK) pathways which explains the mechanisms behind inhibition of TGFβ induced hPSC activation. Furthermore, inhibition of the gene expression of direct ITGA5-related signaling molecules such as FLH3 and Paxillin, as well as downstream factors (CDC42, RAC1 and RHOA) in ITGA5 knockdown hPSCs explains the inhibitory effects on migration, proliferation, adhesion and contraction.

Our data show a modulation of the secretory phenotype of hPSCs after ITGA5 knockdown which abrogated hPSC-mediated activation of tumor cells in terms of migration and growth. These findings corroborate with findings of a recent study showing that inhibition of kindlin-2, an integrin-activating focal adhesion protein, leads to inhibition of PSCs-induced tumor cell activation ^43^. The hPSC-induced fibrosis and tumor-promoting effects were confirmed *in vivo* in co-injection tumor model, as shown with induced fibrosis markers (α-SMA and Collagen I) and tumor progression. These findings are consistent with previous studies showing PSC-induced pancreatic tumor growth ^11, 12^. In line with our *in vitro* data, ITGA5 knockdown in hPSCs led to reduced fibrosis in tumors which resulted in a reduced tumor progression. To confirm that the reduced tumor progression was not due to the death of hPSCs, we confirmed the presence of activated hPSCs using α-SMA immunostaining.

Furthermore, our novel AV3 peptidomimetic against ITGA5 robustly showed a specific binding to ITGA5 receptor resulting in reduced TGFβ-mediated activation of hPSCs by inhibiting FAK pathway. To emphasize, we demonstrate that TGFβ has a crosstalk with ITGA5 and thereby activates integrin-mediated FAK pathway (**Fig. 6G**). Interestingly, blocking of ITGA5 with AV3 inhibited the activation of FAK pathway, suggesting that AV3 likely interacts with ITGA5 at a specific site which is involved in the crosstalk with TGFβ. Therefore, treatment with AV3 inhibited TGFβ-induced collagen and α-SMA in hPSCs. *In vivo*, treatment with AV3 either systemically (i.p) or locally (i.t.) reduced the tumor growth due to reduction of fibrosis, which is in line of the sh-ITGA5 hPSCs *in vivo* data. It is well established that tumor stroma acts as a barrier for chemotherapy penetration. Remarkably, in a PDX tumor model for pancreatic tumor, the combination of AV3 with gemcitabine potentiated the anti-tumor effects leading to >80% reduction in tumor growth. Histologically, we proved that the effects were related to reduction in fibrosis leading to decompression and normalization of blood vessels which allows a better tumor drug perfusion, as shown with the ICG imaging experiment. Surprisingly, despite the reduction in fibrosis after the treatment with AV3, there was no direct effect on the tumor growth. This might be due to replacement of patient CAFs by mouse fibroblasts, which likely have different phenotype and do not contribute to the tumor growth.

In conclusion, this study reveals ITGA5 as a novel prognostic and therapeutic target in pancreatic cancer with a strong impact on the regulation of PSC-induced desmoplasia in pancreatic tumor. In addition, it sheds new light on the importance of TGFβ-mediated activation of ITGA5-FAK signaling in PSCs. The new endogenous peptidomimetic AV3, derived from fibronectin showed blockade of the latter signaling pathway, which led to inhibition of TGFβ-mediated PSC activation. Most excitingly, co-treatment of the peptidomimetic with chemotherapy enhanced the efficacy of chemotherapy in a PDX model due to reduction in tumor fibrosis. These findings are highly promising and suggest a relatively simple translation into the clinic for the treatment of pancreatic cancer.

## Materials and Methods

### Study design

This study was designed to show ITGA5 as a therapeutic target in PSCs and to assess the therapeutic potential of the novel peptidomemtic (AV3) in combination with chemotherapy in PDx pancreatic tumor model. We investigated the prognostic value of ITGA5 in PDAC patients and the tissue samples were obtained during 2001 to 2012 from 137 patients with pancreatic adenocarcinoma. *In vitro* assays (qPCR, Western blot, immunocytochemistry, migration assay, contraction assays) and *in vivo* xenograft tumor models (co-injection tumor model and patient-derived xenograft models) were used to investigate ITGA5 as a therapeutic target in PSCs and the efficacy of novel peptidomimetic AV3. The majority of findings were evaluated by more than one method, and experiments were repeated multiple times independently. *In vivo* experiments were randomized and partly blinded and the number of samples were calculated using Power test considering meaningful differences, % coefficient of variation within the experiment and minimum p-value of 0.05. Cultured cells or PDx tissues were used for these animal models. All experiments conducted under the international animal ethical guidelines, which were approved by the animal ethical committee of Utrecht University, The Netherlands.

### Cells

Human pancreatic stellate cells (hPSCs) were purchased from ScienCell (Carlsbad, CA) and were maintained in a special culture medium provided by the manufacturer, supplemented with 1% penicillin/streptomycin. Pancreatic cancer cell line (PANC-1) was obtained from the American type culture collection (ATTC, Rockville, MD) and, was cultured in Dulbecco’s modified Eagle’s medium (DMEM, PAA, The Netherlands) supplemented with FBS (10%) and antibiotics (1% penicillin- streptomycin). To knockdown ITGA5, hPSCs were transfected with custom-made lentiviral vector shRNA plasmids against ITGA5 (Gene ID: 3678) under puromycin resistance (ATCGbio Life Technology Inc., Burnaby, Canada) using Lipofectamine 2000 transfection reagent (Thermo Fisher Scientific, Breda, the Netherlands).

### Design of AV3 peptidomimetics

To select a peptidomimetic ligand against ITGA5, overlapping sequences (12 aa. long with 8 aa. overlaps) from human FN-III domains-9, and 10 were designed and displayed on a cellular membrane. The domains 9 and 10 of FN were chosen to design peptidomimetics, as these domains were reported to be responsible for binding to the α5β1 receptor, as shown with the docking experiments ^31^. The interaction studies were performed against human recombinant integrin α5β1 receptor (R&D systems) and the bound proteins were transferred to another membrane, and ITGA5 was detected with antibodies. AV3 (Arg-Tyr-Tyr-Arg-Ile-Thr-Tyr) and AV3-PEG6-5FAM (AV3-FAM) were custom-synthesized by China Peptide Co., Ltd. (Shanghai, China). The %purities were 98% for AV3 and 95% for AV3-PEG_6_-5FAM, as assessed by reversed-phase high-performance liquid chromatography (HPLC, analytical). The products were stored at −20°C.

### Patient material, immunohistochemistry and scoring method

Retrospectively collected, formalin-fixed and paraffin-embedded tissue blocks were obtained from the archives of the pathology department for 137 patients with pancreatic adenocarcinoma, who underwent resection with curative intent during the period from 2001 to 2012 at the Leiden University Medical Centre, Leiden, The Netherlands. Only patients with pancreatic adenocarcinoma were included in this study. None of the patients received chemotherapy and/or radiation prior to surgery. Clinicopathological data were collected from electronic hospital records. Differentiation grade was determined according to the guideline of the World Health Organization, and the TNM stage was defined according to the American Joint Commission on Cancer criteria. All samples were non-identifiable and used in accordance with the code for proper secondary use of human tissue as prescribed by the Dutch Federation of Medical Scientific Societies. The use of archived human tissue was conformed to an informed protocol that had been reviewed and approved by the institutional review board of the Leiden University Medical Centre, Leiden, The Netherlands.

Tissue microarrays were prepared to examine the expression of ITGA5 and α-SMA. For each patient triplicate 2.0 mm cores were punched from areas with clear histopathological tumor representation (determined by hematoxylin-eosin staining) from formalin-fixed-paraffin-embedded tissue blocks of their primary tumor and transferred to a recipient TMA block using the TMA master (3DHISTECH, Budapest, Hungary). From each completed TMA, 5 μm sections were sliced and deparaffinized in xylene, rehydrated in serially diluted alcohol solutions, followed by demineralized water. Endogenous peroxidase was blocked by incubation in 0.3% hydrogen peroxide in phosphate-buffered saline (PBS) for 20 min. Antigen retrieval was performed by heat induction at 95°C using citrate buffer (pH 6.0, Dako, Glastrup, Denmark). Tissue microarray sections were incubated overnight with antibodies against ITGA5 (HPA 002642; Sigma-Aldrich®) and α-SMA (PA5-16697; Thermo Fisher Scientific). Negative control samples were incubated with PBS instead of the primary antibodies. The slides were then incubated with Envision anti-rabbit (K4003; Dako) for 30 min at room temperature. After additional washing, immunohistochemical staining was visualised using 3.3-diaminobenzidine tetrahydrochloride solution (Dako) for 5-10 min resulting in brown color and then counterstained with hematoxylin, dehydrated and finally mounted in pertex. All stained sections were scanned and viewed at 40% magnification using the Philips Ultra-Fast Scanner 1.6 RA (Philips, Eindhoven, The Netherlands).

Tumor/stroma ratio was determined on sections stained with haematoxylin and eosin (H&E). The cut-off point for stromal high tumors was the presence of one tumor area with > 50% tumor-stroma. α-SMA staining was scored, according to the extent of stromal positivity, as positive when >50% stroma stained positive. Stromal ITGA5 staining was categorized by multiplying the percentage of stained cells (P) by the intensity of staining (I). Percentage of stained cells: 0 (absence of stained cells), 1 (<25% stained cells), 2 (26-50% stained cells), 3 (>50% stained cells). Staining intensity: 1 (mild), 2 (moderate), 3 (intense). ITGA5 was considered to be positive when the final score (P × I) was > 4.

### Transcriptomics analysis

ITGA5 mRNA expression was analysed in the human cohort from the public database of human pancreatic gene expression datasets from the Expression Omnibus database (GEO). We used GSE28735 dataset consists of pancreatic tumor and adjacent non-tumor tissues from 45 patients with PDAC. The expression levels for ITGA5 from this dataset were statistically compared using two-sided unpaired students’ ttest.

### RNA isolation, reverse transcription and qPCR

Sh-NC or sh-ITGA5 hPSCs were lysed with lysis buffer for total RNA. RNA isolation, cDNA and qPCR performed as described previously ^13^.

### Western blot analyses and immunofluorescent staining

Cells were lysed with SDS-lysis buffer and lysate was loaded on pre-casted Tris-Glycine (4-20% or 10%) gel (Thermo Fisher Scientific, Breda, the Netherlands) and transferred onto PVDF membranes (Thermo Fisher Scientific). The blots were probed with the primary antibodies at different dilutions and were incubated overnight at 4°C, followed by incubation at RT for 1 h with species-specific horseradish peroxidase (HRP) conjugated secondary antibodies. The proteins were detected by the Pierce™ ECL Plus Western Blotting Substrate kit (Thermo Scientific) and exposed to FluorChem™ M System (ProteinSimple, CA). The protein levels were normalized with β-actin and quantified by Image J Software (NIH, MD).

For immunofluorescent staining, cells cultured on 24 well plate were fixed for 20 mins in 4% paraformaldehyde. Then, the cells were incubated with primary antibody and Alexa 488/594 labelled secondary antibodies (Thermo Fisher Scientific). Nuclei were detected with DAPI (4,6-diamidino-2-phenylindole) media and visualized using EVOS fluorescence microscopy (Thermo Fisher Scientific).

### Adhesion assay

Fibronectin (FN, Sigma) was coated in a final concentration of 10μg/ml on 48 well plate at 37°C overnight. Unbound FN was removed with PBS washing. Unspecific binding sites were blocked with 1% BSA for 1 hr at RT. Consequently, sh-ITGA5 and sh-NC hPSCs were seeded (3 × 10^4^ cells/well) and allowed to adhere to the FN coated plates for 30 mins. Unattached cells were removed by PBS washing and adherent cells were fixed in 4% paraformaldehyde and stained with phalloidin labelled with TRITC and DAPI. Phalloidin labelled cells were imaged and counted.

### Spheroid formation assay

Spheroids containing either sh-NC hPSCs or sh-ITGA5 hPSCs were prepared using the hanging drop method as described elsewhere ^13^. hPSCs were suspended in culture medium to a concentration of 2.5 × 10^5^ cells/ml. Approximately five drops (20 μl/drop containing 5 × 10^3^ cells) were distributed onto a lid of a cell culture dish. Then, the lid was inverted and placed over the dish containing PBS for humidity. The spheroids were grown for six days and imaged under an inverted microscope. The diameter of the spheroids was measured digitally using ImageJ software.

### Migration assay

For the scratch assay, sh-ITGA5 and sh-NC hPSCs were seeded in a 24-well plate (6 × 10^4^ cells/well) and allowed to become confluent. A standardized scratch was made using a 200 μl pipette tip fixed in a custom-made holder. Then, cells were washed and incubated in fresh serum free media without growth factors. Images were captured at t = 0 h and t = 15 h, under an inverted microscope. Images were analysed by Image J software to calculate the area of the scratch and represented as the percentage of wound closure compared to control cells.

The transwell migration, assay was carried out in 24 well modified Boyden chamber kit (8μm pores, Corning incorporation, Corning, NY, USA). 5 × 10^4^ PANC-1 tumor cells were seeded in serum-free medium in the inserts with conditioned media from sh-NC or sh-ITGA5 either activated with or without TGFβ as a chemoattractant. After 16-hour incubation, cells migrated on the underside of the membrane were fixed in ice cold 100% methanol and stained with 0.1% crystal violet (Sigma). Migrated cells were counted in 4 random fields at x 100 magnification.

### Cell proliferation assay

Sh-ITGA5 and sh-NC hPSCs proliferation was analysed with a BrdU assay (Roche life sciences, Indianapolis, USA). Cells were plated at a density of 2.5 × 10^3^ cells/well in 96 well-plate. Cells were labelled using 10μM BrdU at 37°C for 2 hours. Cells were fixed by adding FixDenat and incubated with anti-BrdU-POD antibody for 90 minutes. Antibody was removed, cells were washed and substrate solution added. The substrate product was quantified by measuring absorbance at 370nm with a reference wavelength of 492nm.

PANC-1 tumor cell growth was assessed by the AlamarBlue assay. Conditioned media were collected from sh-NC or sh-ITGA5 PSCs with or without activation of TGFβ. PANC-1 cells were seeded at a density of 2.5 × 10^3^ cells/well in a 96 well-plate and treated with the conditioned media obtained from hPSCs. After 3 days of treatment, cells were incubated with AlamarBlue (Invitrogen) at 37°C for 4 hours. Later, fluorescence reading (540 nm excitation and 590 nm emission wavelength) was recorded with VIKTOR™ (PerkinElmer, Waltham, MA).

### 3D collagen I gel contraction assay

A collagen suspension (5 ml) containing 3.0 ml Collagen G1 (5 mg/ml, Matrix biosciences, Morlenbach, Germany), 0.5 ml 10x M199 medium (Sigma), 85 ul 1N NaOH (Sigma) and sterile water was mixed either with 1.0 ml (2 × 10^6^ cells) sh-ITGA5 or sh-NC hPSCs. Collagen gel-cells suspension (0.6 ml/well) was plated in a 24-well culture plate and allowed to polymerize for 1 h at 37 °C. Once polymerized, 1 ml of serum free medium was added with or without TGFβ (5 ng/ml) followed by detachment of the gels from the culture wells. To study the effect of peptidomimetic AV3, 1 ml of serum free medium with TGFβ (5 ng/ml) and 20μM AV3 or sAV3 peptidomimetic was added to the detached gels. Representative images were made at either at 72 h or 96 h using a digital camera (Nikon, Mississauga, ON, Canada). Measurement of collagen gel diameter was performed using Image J imaging software (NIH, Bethesda, MD).

### Cell binding assay

hPSCs or ITGA5 KD hPSCs were seeded in 96 well plate at a density of 2.5 × 10^3^ cells/well. Next day, cells were starved and activated either with or without TGFβ. After 24 h, cells were washed with PBS and fixed with 4% formaldehyde (Sigma) for 15 min at room temperature. Later, fixed cells were blocked in serum free medium with 1% BSA for 30 minutes at room temperature. The cells were incubated with 10μM AV3-FAM in serum-free medium with 0.1% BSA for 60 mins at room temperature. After incubation, the cells were washed with PBS, the nuclei stained with DAPI followed by imaging using EVOS fluorescence microscope (Thermo Fisher Scientific).

### Flow cytometry

hPSCs were seeded at a density of 4 × 10^5^ cells per T25 flask. Next day, cells were starved and activated either with or without TGFβ. After 24h, cells were trypsinized, and cell numbers were diluted to 1 × 10^5^ cells/ml. Cells were incubated at 37°C for 30 min to allow receptor recovery. Then different concentration (1, 2.5, 5, 10μM) of AV3-FAM were added to the suspension cells containing 2% FBS and incubated at 4°C for an hour. Cells were then centrifuged at 300g at 4°C for 5 min. The supernatant was decanted without disturbing the pellet and cells were washed 3 times with 0.5% FBS cold PBS and then fixed in 0.5% formaldehyde for 10 min at 4°C. Cell fluorescence was measured with flow cytometry (BD FACSCalibur™).

### Animal experiments and ethics statements

All the animal experiments in this study were performed with the guidelines, and approved by the animal ethical committee of Utrecht University (2014.III.02.022), the Netherlands. Collection of patient tissue material was approved by the AMC ethical committee (BTC 2014_181), and performed according to the Helsinki Convention guidelines. Informed consent was obtained for all inclusions. Grafting of immune deficient NSG mice with patient material was performed according to procedures approved by the animal experiment ethical committee (DTB102348/LEX268).

### Pancreatic co-injection xenograft and patient-derived xenografts (PDX) models

Six-week-old male CB17 SCID mice (Janvier labs) were subcutaneously (s.c.) injected with PANC-1 alone (2 × 10^6^ cells) or co-injected with hPSCs (2 × 10^6^ cells) or stably transfected with either sh-ITGA5 or sh-NC hPSCs (2 × 10^6^ cells) followed by tumor measurement every 2-3 days.

For AV3 peptidomimetic study, animals were subcutaneously co-injected with PANC-1 (2 × 10^6^ cells) and hPSCs (4 × 10^6^ cells). Six tumor-bearing mice per treatment groups were taken and injected with either vehicle, AV3, or sAV3. Half of the group was injected intra-peritoneal (20mg/kg) and the other 3 intra-tumoral (4mg/kg). Injections were given twice a week starting from day 9.

Freshly excised pancreatic patient tumor piece grafted subcutaneously into the flank of immunocompromised NOD.Cg-Prkdc^scid^Il2rg^tm1Wjl^/SzJ (NSG) mice with matrigel as passage 1. After transplantation, tumor growth was monitored, and upon reaching a size of 800-1000mm^3^, PDX tumors were harvested and transplanted into the flank of NOD/SCID animals with Matrigel as passage 2. After that, mice with tumors reaching around 150mm3 were injected intraperitoneally with either vehicle, 20mg/kg of AV3, 50mg/kg of gemcitabine or AV3 (20mg/kg) and Gemcitabine (50mg/kg) twice a week for 3 weeks (n=5 mice per group). To study the effect of AV3 on the tumor perfusion, indocyanine green (5mg/kg) was injected via tail vein. After 24 hrs of the injection, tumors were imaged using a small animal imager (Pearl imager, LICOR, Lincoln, NE).

Tumor growth was assessed with calliper every 2-3 days. Tumor volumes were measured by the formula (V=L × B2/2). At the end of the experiments, animals were sacrificed under anaesthesia after which tumors were harvested and immediately snap frozen in cold 2-methyl butane. Frozen organs were stored at −80°C until analysis. Cryosections (4μm) were cut and fixed in acetone for 10 mins before staining for collagen-1, and αSMA (Table 2) following by fluorescent secondary antibody and the protocol described elsewhere ^44^.

### Statistical analyses

Regarding survival analyses: Overall survival (OS) was defined as the time from the date of surgery to the date of death or lost to follow-up. Kaplan-Meier estimates of the survival function, including P values from the log-rank test were used to graphically compare the time-to-event outcomes based on ITGA5 expression and to estimate median OS. Furthermore, univariate and multivariate survival analyses were performed using the Cox proportional hazard regression model. Next to age and gender only variables that were significant in univariate analysis were included in multivariate analyses, with an exception for pT-stage for OS (SPSS statistical software, version 23.0; IBM SPSS Inc, Chicago, IL). Group data presented as mean ± SEM for at least three independent experiments. The graphs and statistical analyses were performed using GraphPad Prism version 5.02 (GraphPad Prism Software, Inc., La Jolla, CA). Statistical analysis of the results was performed either by a two-tailed unpaired student’s t-test for comparison of two treatment groups or a one-way ANOVA to compare multiple treatment groups. In all cases, differences were considered significant at p < 0.05.

## Acknowledgements

Authors thank Ms. Cynthia Waasdorp from Academic Medical Centre, The Netherlands and Ms. Bettie Klomphaar (University of Twente) for their technical assistance in animal experiments.

## Author contribution

P.R.K designed, performed, analyzed and interpreted the experimental data and wrote the manuscript. R.B assisted in *in vivo* experiments and data analysis and critically reviewed, edited the manuscript. J.S supported during the *in vivo* experiments. G.S read the manuscript. M.F.B and H.W.L provided the PDx pancreatic tumor sample and gave inputs to the manuscript. S.W.G, P.J.K, A.L.V, C.F.M.S contributed to the clinical patient data, interpreted and analyzed the data. J.P designed the study, interpreted the studies, wrote and edited the manuscript.

## Supplementary figures and tables

**Fig. S1:**
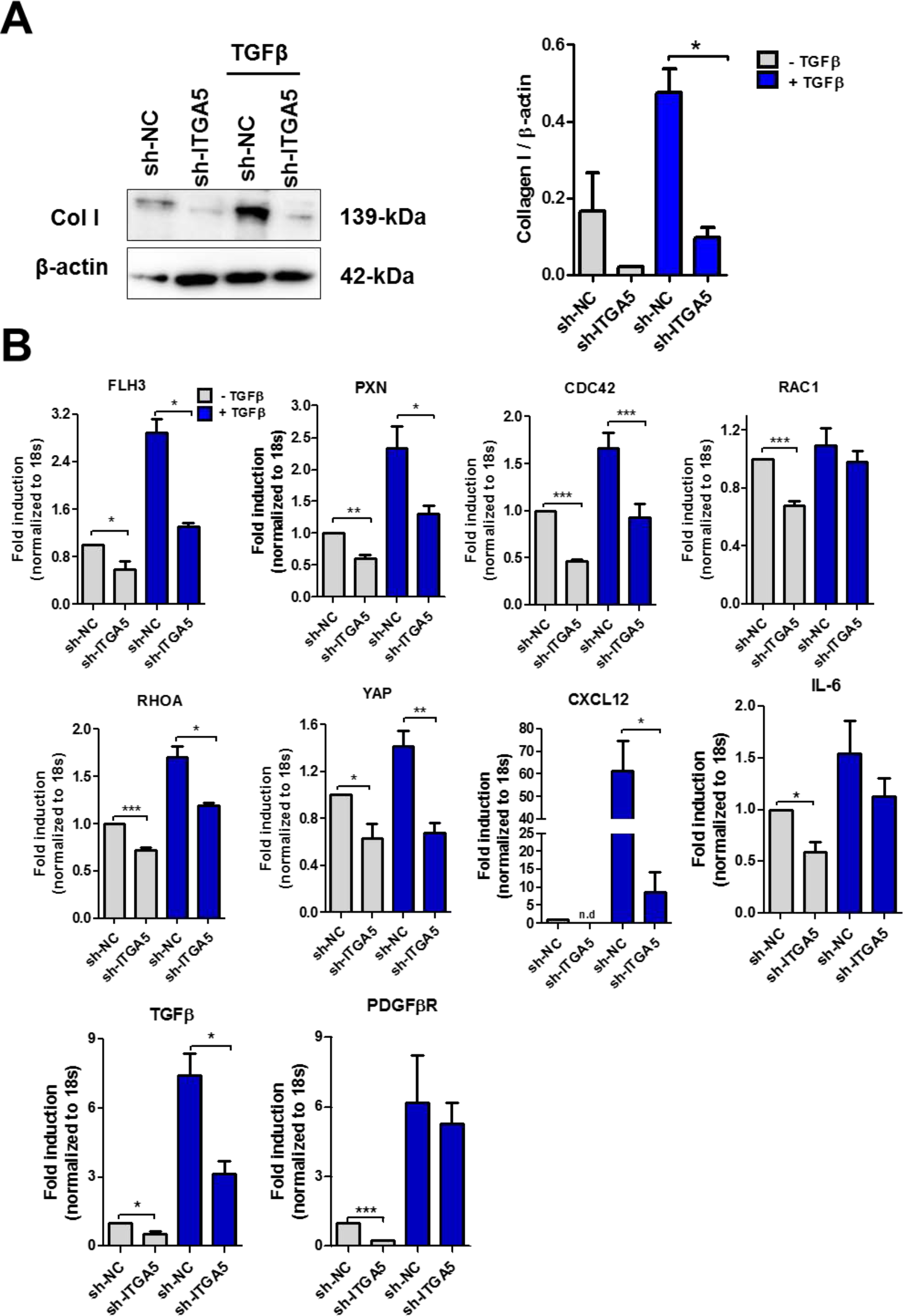
Effect of ITGA5 knockdown in hPSCs on collagen-1 expression and downstream genes. (**A**) Western blot analysis of collagen 1 expression in sh-ITGA5 and sh-NC hPSCs with or without TGFβ activation. (**B**) Gene expression analysis in sh-ITGA5 hPSCs. Data represent mean ± s.e.m from at least three independent experiments; *p<0.05, **p<0.01, ***p<0.001.

**Fig. S2:**
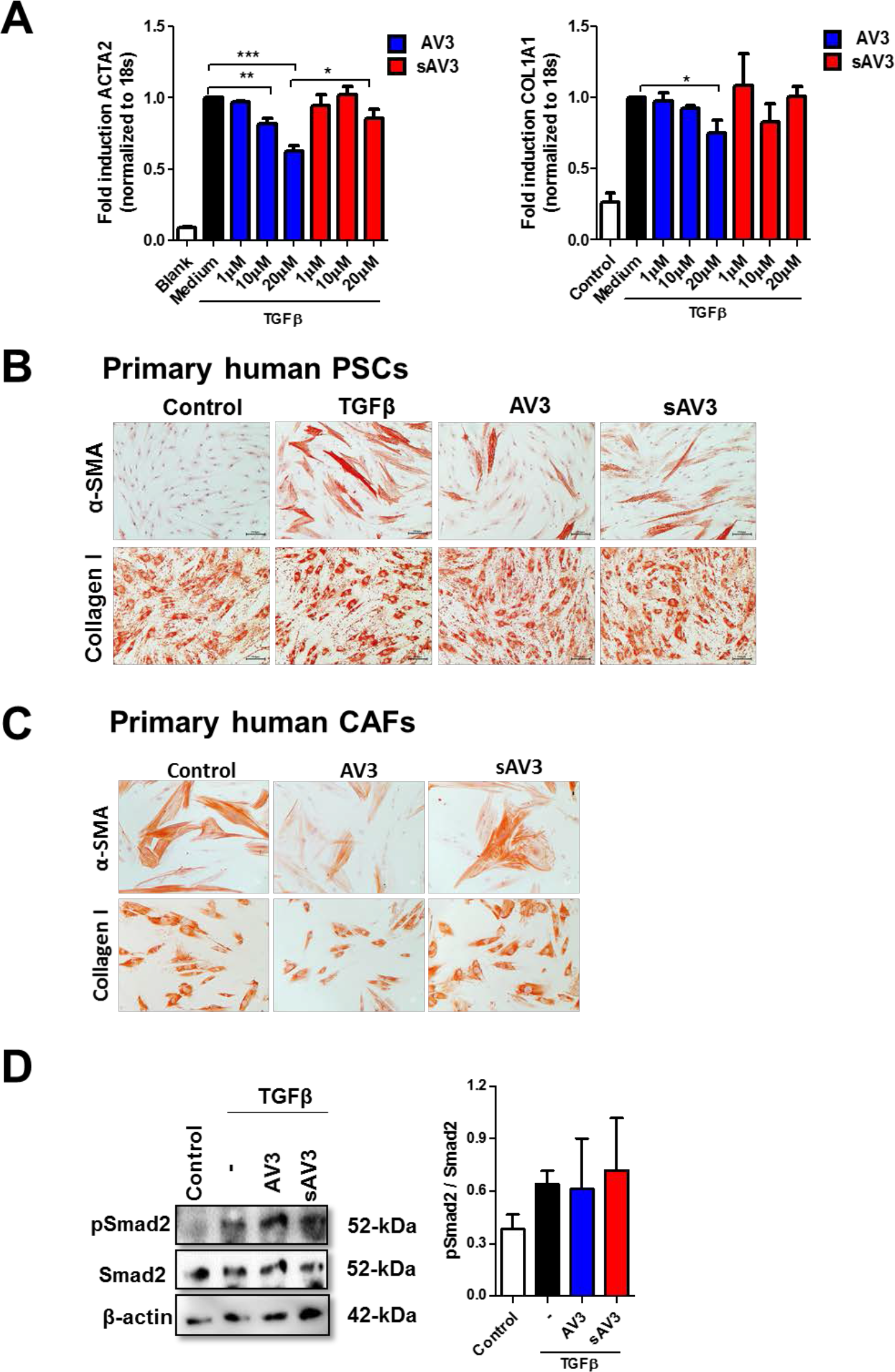
Effect of ITGA5 blocking peptidomimetic (AV3) on hPSCs and human pancreatic CAFs. (**A**) AV3 reduces TGFβ-induced differentiation markers (Acta2 and collagen 1) expression levels in dose dependent manner in hPSCs. (**B, C**) Treatment of primary hPSCs and patient-derived pancreatic primary CAFs with AV3 (20 μM) for 48 hours inhibits differentiation markers such as αSMA and collagen-1 as shown by immunocytochemical staining. Scale bar, 200μm. (**D**) Western blot analyses shows no effect on TGFβ mediated Smad signaling pathway following AV3 treatment in 8 hr TGFβ activated hPSCs. Data represent mean ± s.e.m from at least three independent experiments; *p<0.05, **p<0.01, ***p<0.001.

**Fig. S3:**
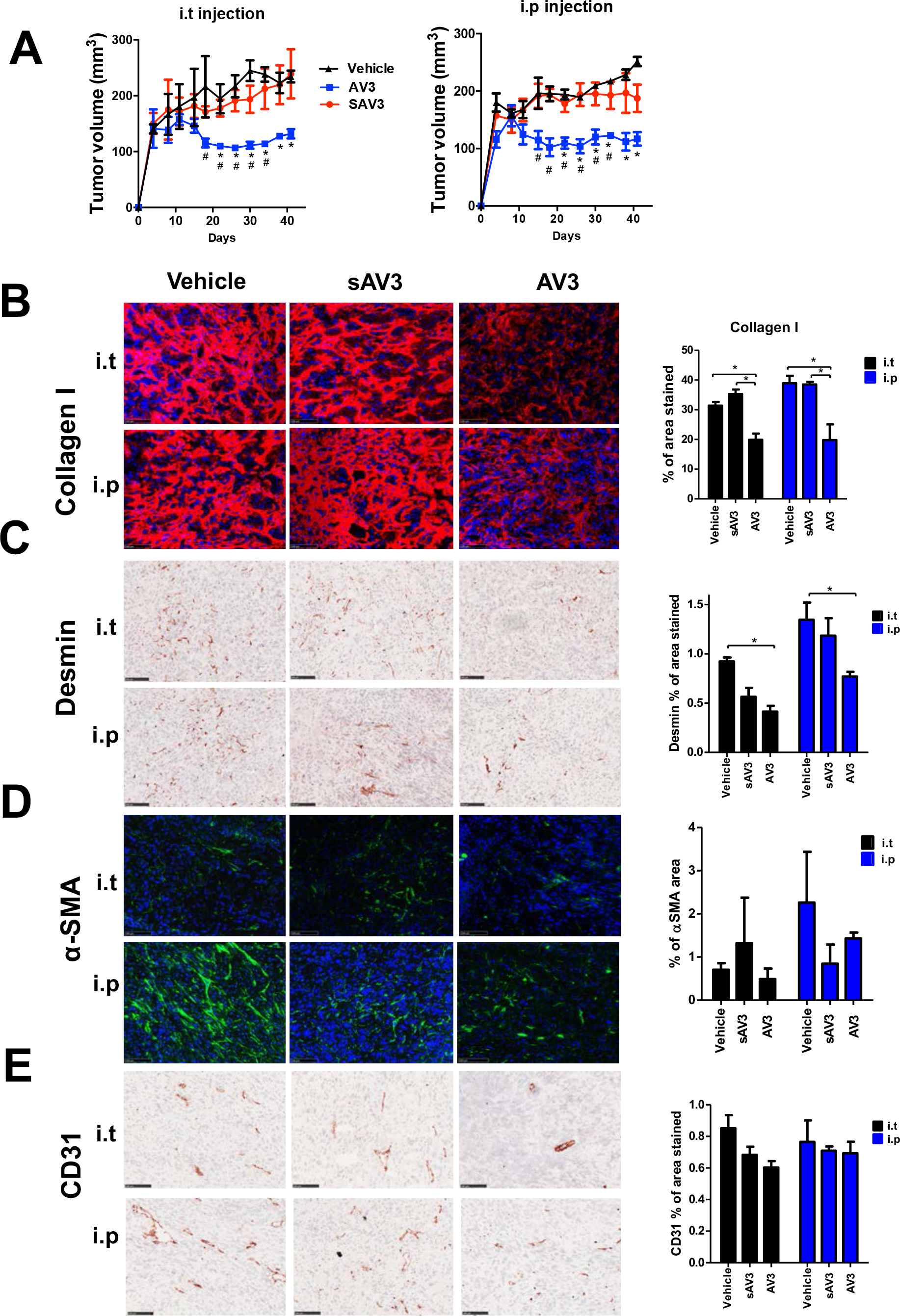
AV3 reduces PSC-induced pancreatic tumor growth in the subcutaneous co-injection (PANC-1+hPSCs) pancreatic tumor model. (**A**) Tumor growth curves demonstrating that administration of AV3 either i.p or i.t. reduces the tumor growth significantly in co-injection xenograft model (PANC-1 + hPSC). (**B-E**) Representative fluorescent microscopic pictures of tumors immunostaining and quantitative analyses showing the effect of AV3 treatment on collagen1, desmin, αSMA and CD31. Data are represented as mean ± s.e.m. *p<0.05 vs. Vehicle; #p<0.05 vs. sAV3.

**Fig. S4:**
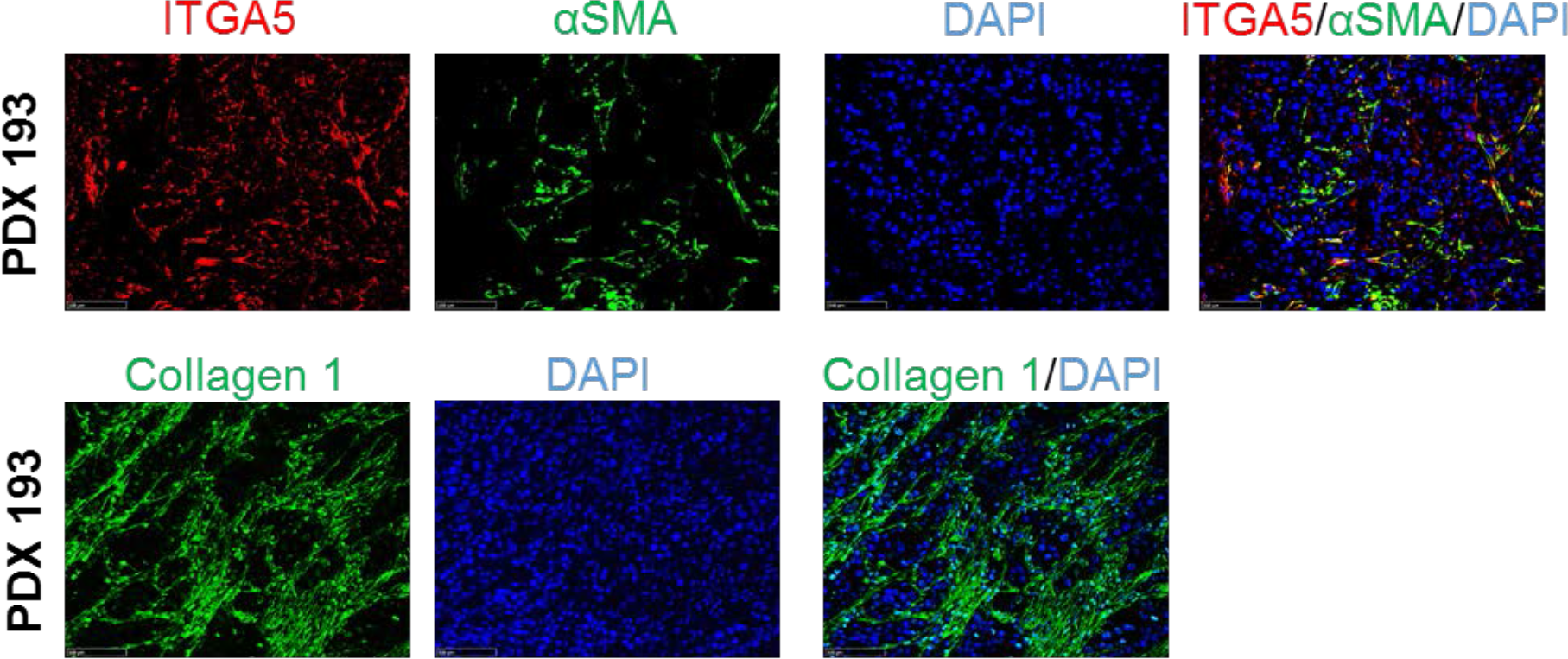
Immunofluoresent staining in pancreatic patient tumor. Immunofluoresence stainings of ITGA5, αSMA, and collagen1 in pancreatic tumor patient (PDX 193). Scale bar, 100μm.

**Fig. S5:**
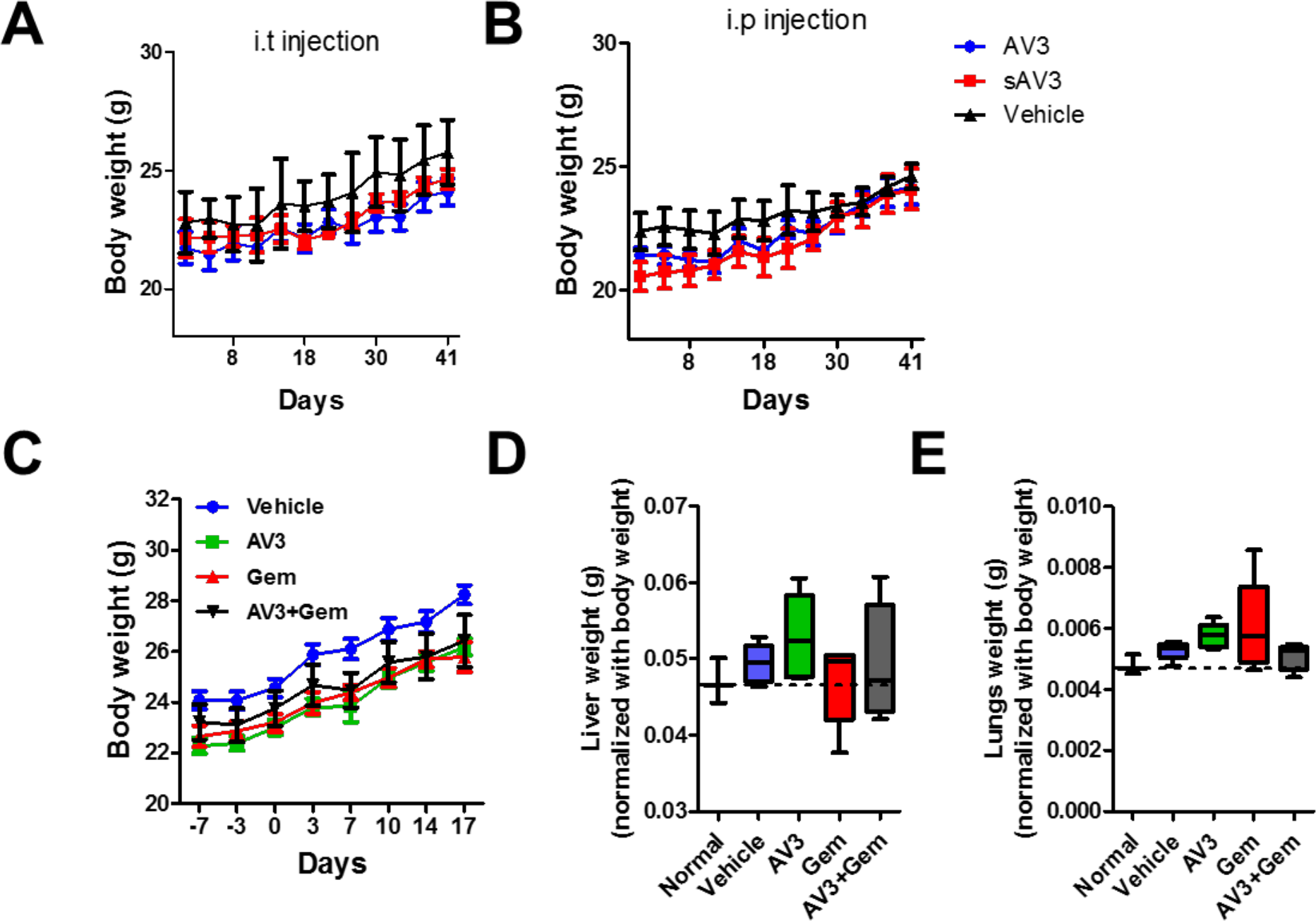
Effect of AV3 on body and organs weight in mice. Body weight showing the effect of different treatments in co-injection tumor model (**A, B**) and PDX tumor model **(C)**. (**D-E**) Graphs show the effect of different treatments on liver and lung weights in PDX model and normal mice. Data represent mean ± s.e.m.

**Table S1:** Characteristics for the pancreatic adenocarcinoma patients.

| **Characteristics** |  |  |
| --- | --- | --- |
| Age, n (%) | <65 years | 66 (48%) |
|  | ≥65 years | 71 (52%) |
| Tumor differentiation, n (%) | Well | 12 (9%) |
|  | Moderately | 43 (31%) |
|  | Poorly | 45 (33%) |
| Tumor stage, n (%) | Stage I | 17 (12%) |
|  | Stage II | 112 (82%) |
|  | Stage III | 8 (6%) |
| Vascular invasion, n (%) | Positive | 45 (33%) |
|  | Negative | 92 (67%) |
| Perineural invasion, n (%) | Positive | 87 (64%) |
|  | Negative | 50 (37%) |

**Table S2:**
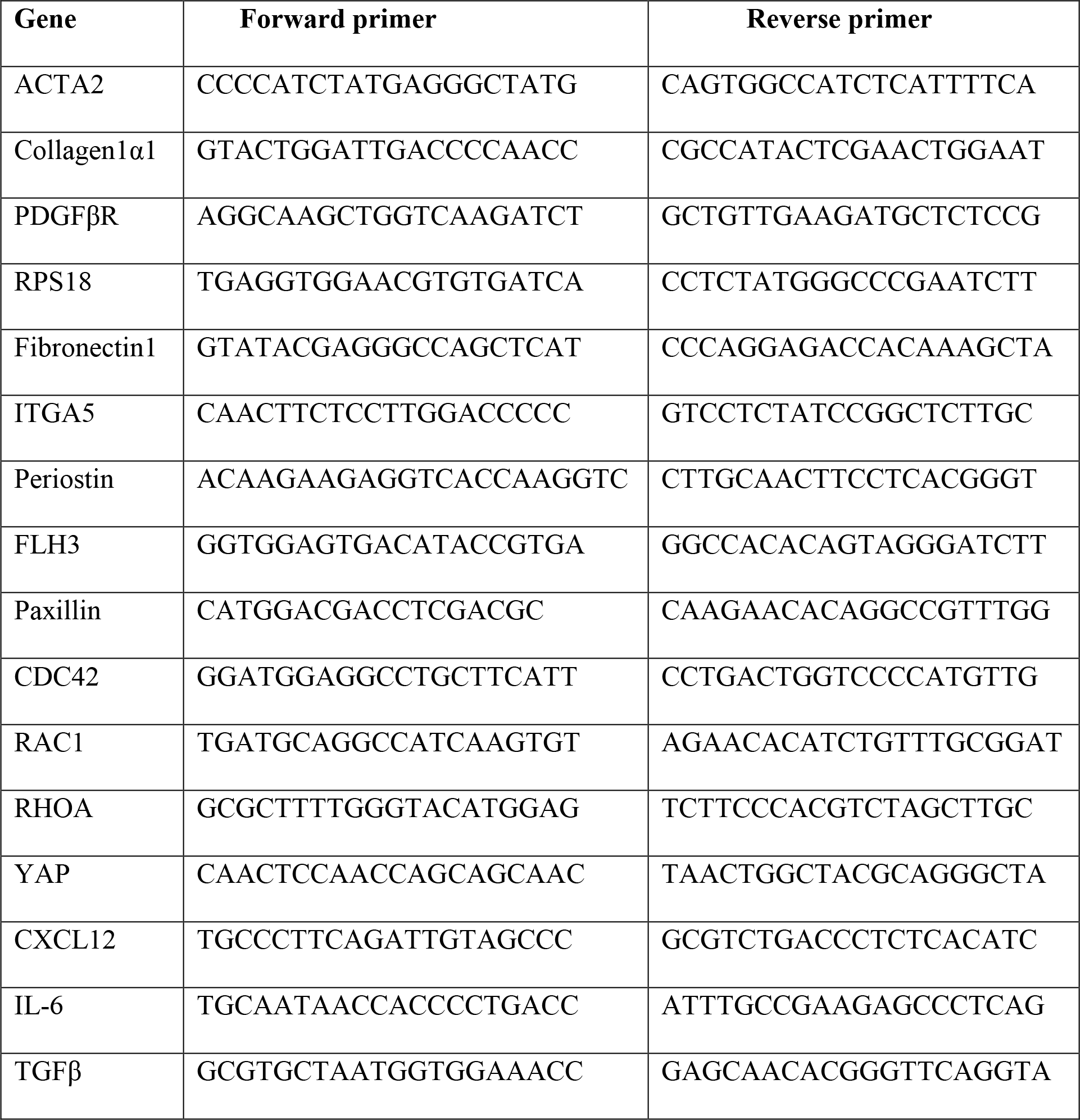
Primers used for quantitative real-time PCR.

**Table S3:** Details of the antibodies used for western blot analyses.

| **Antibody** | **Source** | **Dilution** |
| --- | --- | --- |
| Rabbit polyclonal ITGA5 | Sigma | 1:1000 |
| Mouse monoclonal α-SMA | Sigma | 1:500 |
| Goat polyclonal Collagen1α1 | Southern biotech | 1:250 |
| Mouse monoclonal β-actin | Sigma | 1:5000 |
| Rabbit monoclonal phosho FAK | Cell signaling Tech | 1:1000 |
| Rabbit monoclonal FAK | Cell signaling Tech | 1:1000 |
| Rabbit monoclonal phosho Smad2 | Cell signaling Tech | 1:1000 |
| Rabbit monoclonal Smad2 | Cell signaling Tech | 1:1000 |
| Rabbit monoclonal phospho Akt | Cell signaling Tech | 1:1000 |
| Rabbit monoclonal Akt | Cell signaling Tech | 1:1000 |
| HRP-conjugated rabbit anti-goat IgG | DAKO | 1:2000 |
| HRP-conjugated goat anti-rabbit IgG | DAKO | 1:2000 |
| HRP-conjugated goat anti-mouse IgG | DAKO | 1:2000 |
| Alexa Flour 594 donkey anti-rabbit IgG | Thermo Fisher Scientific | 1:200 |
| Alexa Flour 488 donkey anti-goat IgG | Thermo Fisher Scientific | 1:200 |

**Data file S1: The complete gene array data on sh-ITGA5 hPSCs versus sh-NC and the effect of TGF-β activation on sh-NC and sh-ITGA5 hPSCs.** The .xls file is attached separately.

